# A pretrained unified model enables cellular functional profile prediction and multi-objective virtual drug screening

**DOI:** 10.64898/2026.08.25.746866

**Authors:** Ruoqiao Chen, Li Huang, Yu Qiao, Saurabh Mandal, Linqing Mo, LingXiao Li, Dmitry Leshchiner, Xiaodan Zhang, Jing Pu, Yuying Xie, Reda Girgis, Edmund Ellsworth, Ling Huang, Xin Chen, Xiaopeng Li, Jiayu Zhou, Bin Chen

**Affiliations:** Department of Pharmacology and Toxicology, College of Human Medicine, Michigan State University, Grand Rapids, MI 49503, USA; Department of Pediatrics and Human Development, College of Human Medicine, Michigan State University, Grand Rapids, MI 49503, USA; Hawaii Cancer Center, University of Hawaii, Honolulu, HI 96813, USA; Pancreatic Cancer Program, Henry Ford Health, Detroit, MI 48202, USA; Department of Computer Science and Engineering, College of Engineering, Michigan State University, East Lansing, MI 48824, USA; School of Information, University of Michigan, Ann Arbor, MI 48109, USA; Department of Computational Mathematics and Engineering, College of Engineering, Michigan State University, East Lansing, MI 48824, USA; Corewell Health, Grand Rapids, MI 49503, USA; Department of Medicine, College of Human Medicine, Michigan State University, East Lansing, MI 48824, USA; Center for AI-Enabled Drug Discovery, Michigan State University, Grand Rapids, MI 49503, USA

## Abstract

Cells are characterized by molecular states, coordinated molecular interactions, regulatory programs, and responses to perturbations. Systematic mapping of these cellular functional profiles across biological contexts remains experimentally costly and fragmented. Here we present InsilicoCell, a pretrained multi-modal, multi-task model that unifies prediction of cellular functional profiles spanning molecular states, molecular interactions, and perturbation-induced responses. Built on a supervised transformer architecture and pretrained on more than 88 million measurements across seven tasks, including drug sensitivity, drug-induced gene expression, and drug-protein binding, InsilicoCell learns a shared representation that links molecular profiles to cellular phenotypes, improves performance over task-specific models, and generalizes to unseen entities, contexts, and conditions. InsilicoCell extends beyond cell line systems to patient, spatial and single-cell settings, and enables multi-objective virtual drug screening. It identifies novel candidate compounds with experimental validation, including c-Myc activity inhibitors, antifibrotic agents and stemness-inducing compounds. Together, InsilicoCell provides a scalable framework for predictive cellular biology and therapeutic discovery.

## Introduction

Mapping cellular functional profiles across diverse biological contexts is central to understanding diseases and discovering therapeutics. Despite advances in high-throughput technologies, comprehensive characterization across modalities and conditions remains fragmented and costly^1–3^. Machine learning offers a scalable alternative, yet existing models are typically restricted to individual tasks, such as modelling drug sensitivity^4–6^, perturbation-induced gene expression^7–11^ or molecular interactions^12–15^, and are confined to the contexts represented in their training data, limiting their ability to generalize and integrate knowledge across diverse biological contexts.

Inspired by recent foundation models in other fields^16–18^, we present InsilicoCell, a pretrained, multi-modal unified model for the prediction of cellular functional profiles spanning molecular states, molecular interactions, and responses to perturbations. InsilicoCell integrates diverse molecular modalities, including drugs, genes, and proteins, together with cellular contexts (encoded by transcriptomic profiles) and experimental conditions such as dose and time. Built on a multi-task transformer architecture and pretrained on more than 88 million measurements across seven tasks including drug sensitivity, drug-induced gene expression, drug-protein binding, transcription factor (TF)-gene regulation, gene effect scores from genetic perturbation screens, gene mutation, and copy number variation (CNV), InsilicoCell learns a shared representation among the tasks. This unified framework improves performance relative to task-specific models, enables prediction across previously unseen entities, biological contexts, and conditions, and can be extended to new tasks such as drug combination synergy prediction through transfer learning. Notably, InsilicoCell generalizes beyond cell line data to patient-level, spatial and single-cell settings through zero-shot inference and few-shot adaptation.

Different from recent efforts in virtual cell modeling^19–24^, InsilicoCell provides a unified framework tailored for in silico drug discovery. Conventional drug discovery workflows often separate target-based and phenotypic discovery into distinct stages, creating a gap between molecular activity and cellular efficacy that contributes to translational inefficiency^25–28^. In contrast, InsilicoCell enables multi-objective screening within a single model by jointly modelling molecular interactions (e.g., drug-protein binding) and cellular responses (e.g., drug sensitivity and drug-induced gene expression changes). Building upon our recently developed GPS framework for transcriptomics-based virtual compound screening^8^, InsilicoCell substantially expands the range of predictable biological contexts and screening objectives. This integrated framework allows early prioritization of candidates that satisfy multiple biological properties, thereby reducing late-stage attrition and improving screening efficiency.

We demonstrate the utility of InsilicoCell through three applications with experimental validation: identification of novel c-Myc activity inhibitors in hepatocellular carcinoma (HCC), discovery of compounds that suppress fibrosis markers in idiopathic pulmonary fibrosis (IPF), and identification of compounds that induce cellular stemness markers. Together, these results establish InsilicoCell as a scalable and generalizable pretrained model for predictive cellular biology and therapeutic discovery.

## Results

### InsilicoCell model

InsilicoCell is a pretrained multi-task, multi-modal model for unified prediction of cellular functional profiles spanning molecular states, molecular interactions and perturbation-induced responses. These profiles comprise seven prediction tasks: (1) drug-protein binding affinity, (2) TF-target gene regulation, (3) gene mutation, (4) CNV, (5) drug sensitivity, (6) drug-induced gene expression change, and (7) gene effect score prediction (Figure 1A-1C, S1A-1C). To pretrain and test InsilicoCell, we integrated curated datasets of the seven types of cellular functional profiles from multiple public databases, including LINCS^1^, BindingDB^29^, ChEA^30^, CTRP^31^, and DepMap^2^ (Table S1). Although drug-protein binding and TF-target gene regulation are not cell context-specific tasks, their large-scale datasets capture complementary biological knowledge that could benefit the learning of other context-dependent tasks through shared representations. The combined data include over 88 million labeled samples, covering diverse molecular modalities (816,230 drugs, 18,545 genes, 6,550 proteins/TFs), cellular contexts (1,204 cell lines), and experimental conditions (five time points, six dosages) (Figure 1B). In addition, we incorporated external datasets including GDSC^32,33^, ChEMBL^34^, and GEOMeta (Methods), to evaluate the three drug discovery-related tasks that constitute a primary focus of this study (Table S1).

**Figure 1:**
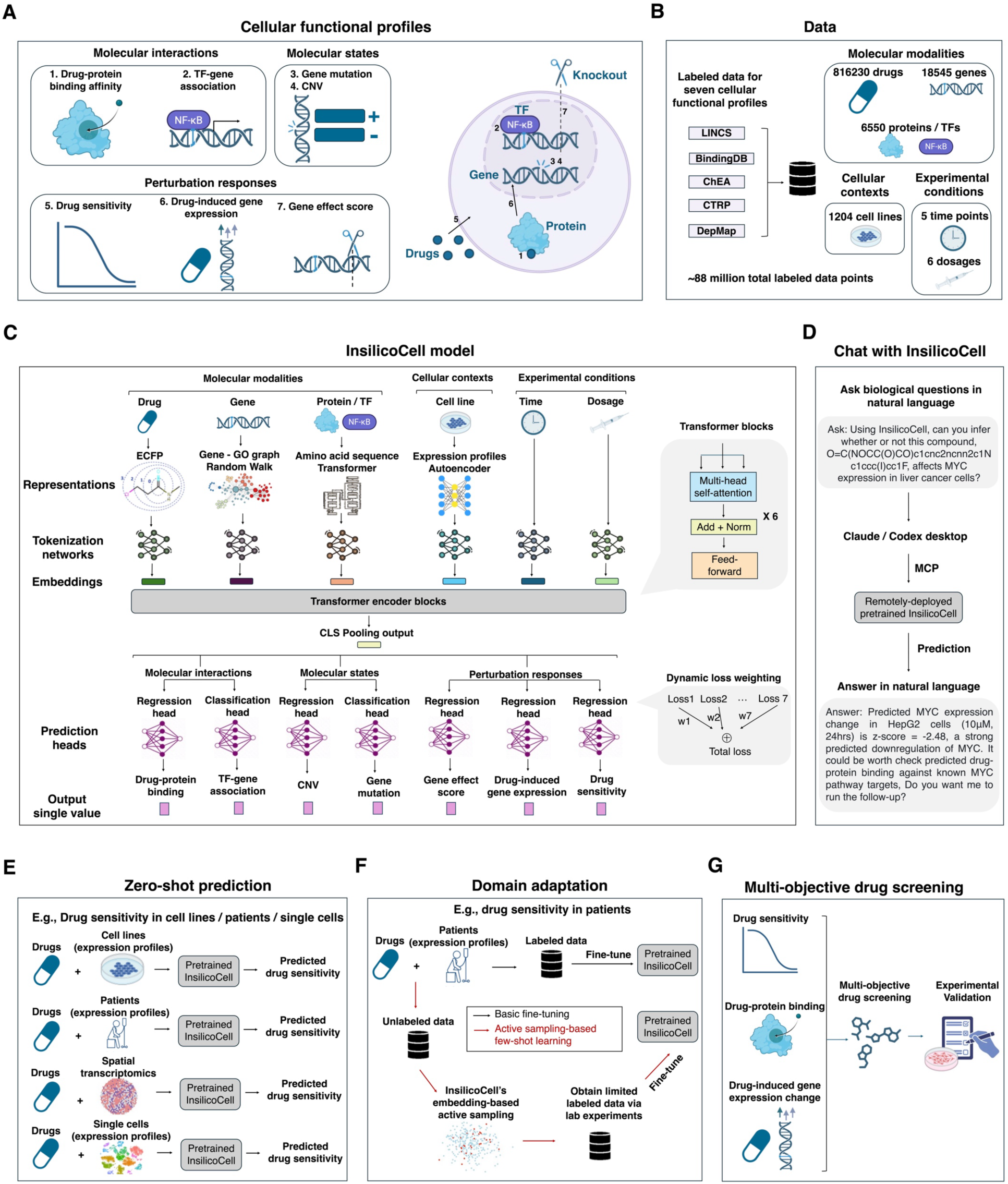
Overview of InsilicoCell. **A**, Unified prediction of cellular functional profiles spanning molecular states, molecular interactions and perturbation-induced responses by InsilicoCell. Seven types of cellular functional profiles are included in the pretraining of InsilicoCell, each formulated as a distinct learning task. **B**, InsilicoCell leverages labeled data from various databases for model pretraining and testing, comprising over 88 million labeled measurements, covering diverse molecular modalities (drugs, genes, and proteins / TFs), cellular contexts (cell lines), and experimental conditions. For drug-induced gene expression prediction, multiple time points and doses are explicitly encoded as model inputs. **C**, Architecture of InsilicoCell. The framework consists of an input representation module involving multi-modal representation and tokenization to generate embeddings of a unified dimension across all inputs, and a multi-task transformer module with task-specific heads for label prediction. A detailed architecture is shown in Figure S1. **D**, Chatting with InsilicoCell. We developed a user-friendly Model Context Protocol (MCP) interface, enabling users to use natural language to interact with InsilicoCell in frontier models including Claude and Codex. **E**, Pretrained InsilicoCell enables zero-shot prediction of the seven cellular functional profiles in various contexts. Shown is an example of predicting in vitro or in vivo drug sensitivity at bulk, single-cell or spatial levels for unseen compounds. **F**, Complementary to zero-shot prediction, InsilicoCell is equipped with domain adaptation approaches to improve performance in new domains. An example is using limited labeled drug sensitivity data in patients to fine-tune InsilicoCell for better patient-level drug sensitivity prediction. An active learning-based framework is adopted by InsilicoCell for improving few-shot adaptation performance under constrained experimental budgets. **G**, InsilicoCell also serves as a virtual platform for multi-objective compound screening. By combining its prediction of multiple types of cellular functional profiles including drug-induced gene expression change, drug sensitivity, and drug-protein binding affinity, InsilicoCell prioritizes compounds which simultaneously satisfy multiple screening objectives for downstream experimental validation.

The architecture of InsilicoCell consists of a multi-modal input representation module, and a multitask transformer label prediction module (Figure 1C, S1A, S1C). Input entities are represented using modality-specific embeddings, including chemical fingerprints for compounds, Gene Ontology-based representations encoded by node2vec for genes, sequence-based representations encoded by ProtTrans for proteins, and transcriptome-based representations encoded by an autoencoder for cellular contexts (Figure S1A). Alternative embedding methods for each modality can be readily incorporated (Methods). A set of tokenization networks further transform all representations into token embeddings of a unified dimension. In the label prediction module, each input sample for each task is represented as a sequence of multi-modal token embeddings (Figure S1B), which are processed by shared transformer encoder blocks via self-attention mechanisms. Task-specific prediction heads subsequently generate outputs for each type of cellular functional profile. InsilicoCell adopts a two-stage supervised pretraining strategy (Methods). In stage 1, data from all seven tasks are jointly used for large-scale multi-task training, with task contributions balanced through a dynamic loss weighting scheme. In stage 2, the model is further specialized for individual tasks by transferring the pretrained weights from stage 1 and continuing training on task-specific datasets. This strategy enables InsilicoCell to first learn shared biological knowledge across tasks and subsequently refine task-specific predictive capabilities.

To facilitate its use, we implement a Model Context Protocol (MCP) server exposing over 20 APIs, enabling frontier models such as Claude and Codex, as well as emerging agentic tools, to interact with InsilicoCell. Notably, users can naturally interact with remotely deployed InsilicoCell to explore biological questions of interest while leveraging the reasoning capabilities of large language models (Figure 1D).

A central design goal of InsilicoCell is generalization. Through its multi-modal representations, the pretrained InsilicoCell allows cellular function prediction involving new compounds, proteins, and genes under new cellular contexts. Beyond cell line-level prediction, InsilicoCell can be extended to support patient, spatial and single-cell-level prediction through either zero-shot inference (Figure 1E) or domain adaptation via fine-tuning (Figure 1F). Specifically, an embedding-based active learning framework further improves adaptation in data-limited settings by efficiently selecting informative samples for measurements.

By leveraging its unified prediction of diverse cellular functional profiles across biological contexts, InsilicoCell enables multi-objective virtual drug discovery (Figure 1G). For instance, by jointly evaluating three tasks of drug sensitivity, drug-induced gene expression changes, and drug-protein binding affinity within a single framework, InsilicoCell prioritizes compounds predicted to inhibit c-Myc activity in HCC cells. Similarly, by integrating predictions from two tasks, the framework identifies candidate compounds that suppress profibrotic programs in myofibroblasts while minimizing toxicity to normal fibroblasts. More broadly, this unified framework enables simultaneous optimization of multiple biological and pharmacological objectives, thereby facilitating efficient candidate prioritization for downstream experimental validation.

### Benchmarking InsilicoCell across tasks

To evaluate the predictive performance of pretrained InsilicoCell, we investigated individual training stages (Figure 2A) and established a de novo benchmarking framework, as no existing model or benchmark simultaneously covers all seven cellular functional profiles addressed in this study (Methods). We considered two scenarios for evaluation: first, sample-level holdout validation, where a subset of labeled input samples from all tasks were randomly selected as test sets; and second, entity-level holdout validation, where a portion of input entities with their corresponding input samples were held out as test sets. The second scenario was designed for evaluating model generalization to previously unseen entities. Five regression tasks were evaluated using Pearson correlation and Root Mean Squared Error (RMSE), while two classification tasks were evaluated using the F1 score and Area Under the Receiver Operating Characteristic (AUROC).

**Figure 2:**
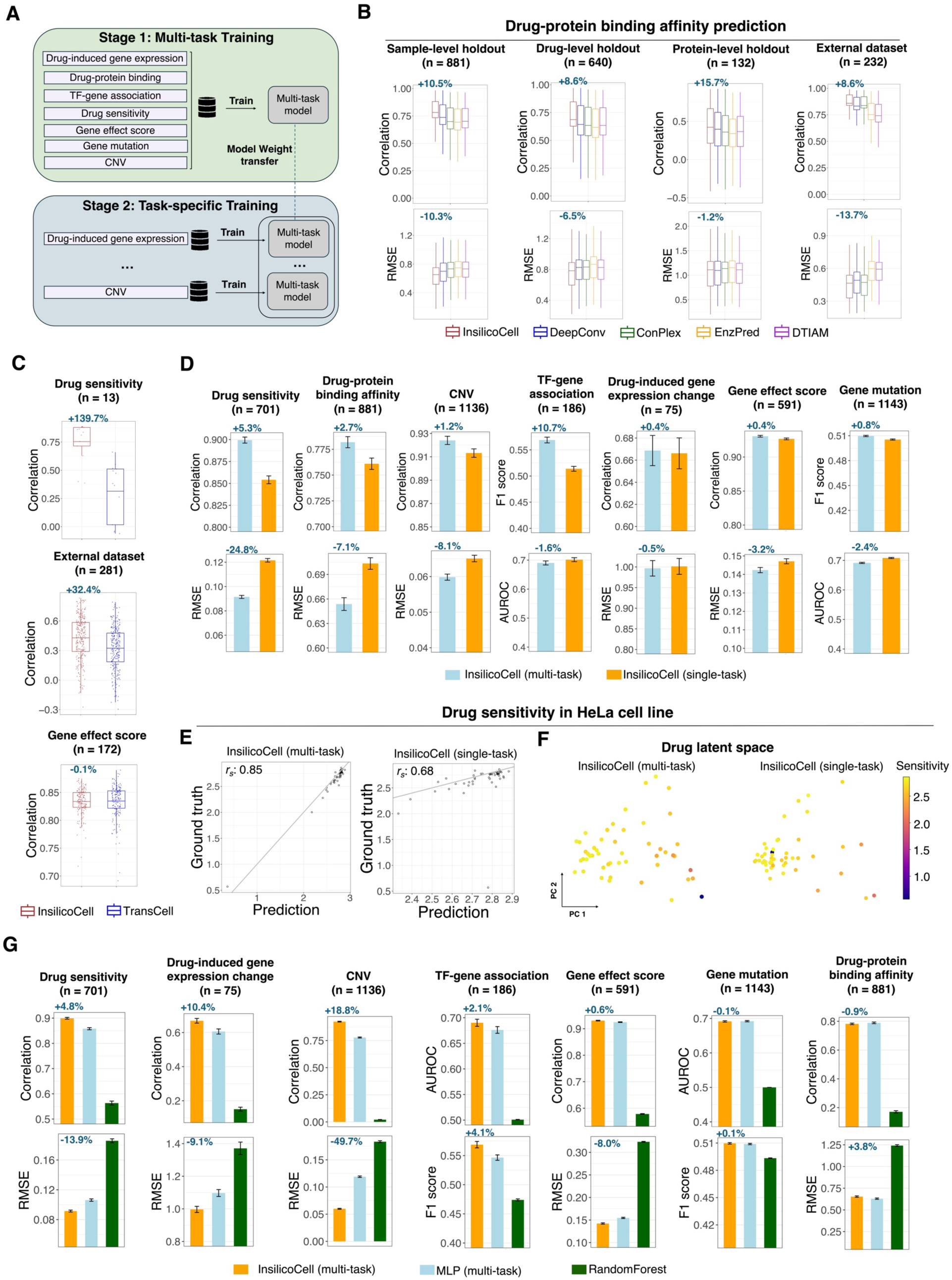
Benchmarking InsilicoCell across cellular functional profiles. **A**, Two-stage training strategy of InsilicoCell. In stage 1, multi-task learning is performed by jointly training the model on data from all tasks. The trained model weights are saved and transferred to stage 2 for task-specific training, where the model is further trained on data for each task separately. **B**, Comparison of prediction performance in drug-protein binding affinity prediction between InsilicoCell and SOTA specialized models. Sample-level, drug-level, and protein-level holdout settings evaluate generalization to unseen samples, compounds, and proteins, respectively. An external dataset was also included for drug-protein binding task evaluation (Methods). Correlation and RMSE were used for evaluation. All the boxplots show the protein-wise performance. *n* denotes the number of proteins. Percentage numbers represent changes in mean protein-wise performance of InsilicoCell compared to the averaged performance among the other four specialized models. **C**, Comparison of drug sensitivity and gene effect score prediction between InsilicoCell and TransCell on unseen cell lines. An external dataset was also included for drug sensitivity task evaluation (Methods). *n* denotes the number of cell lines. Percentage numbers represent changes in mean cell line-wise performance of InsilicoCell compared to TransCell. **D**, Comparison of the prediction performance of multi-task InsilicoCell (two-stage training by default) and single-task InsilicoCell (trained on the data of only a single task from scratch) on the sample-level holdout test set. For drug-protein binding affinity and TF-gene association tasks, the error bars show protein/TF-wise performance (*n* = the number of proteins). For all the other tasks, the error bars show cell-wise performance (*n* = the number of cell lines). All five regression tasks were evaluated using correlation and RMSE and the other two classification tasks were evaluated using F1 score and AUROC. All error bars use standard error of the mean (SE). **E-F**, Representative example illustrating the benefit of multi-task learning in drug sensitivity prediction. **E**, Scatter plots showing the prediction and ground truth for all drugs in the HeLa cell line in the test set. **F**, Visualization of drug embeddings derived from the last hidden layer of the encoder. The color represents the magnitude of the observed drug sensitivity. **G**, Ablation study by replacing the transformer module with baseline architectures including MLP and random forest. Error bars show the prediction performance on the sample-level holdout test set. All error bars use standard error of the mean (SE). Percentage numbers represent changes in mean entity-wise performance of InsilicoCell compared to MLP.

We first compared InsilicoCell with state-of-the-art (SOTA) task-specific models across representative cellular functional profiles. For drug-protein binding affinity prediction, InsilicoCell consistently outperformed established models including DTIAM, DeepConv, ConPlex, and EnzPred^12–15^ across sample-level, drug-level, and protein-level holdout validation settings (Figure 2B, Methods). We next compared InsilicoCell with TransCell^4^, a pretrained model developed for predicting drug sensitivity and gene effect score. To avoid data leakage arising from overlap between InsilicoCell test sets and TransCell pretraining data, we restricted evaluation to cell lines unseen by both models. InsilicoCell outperformed TransCell with a 139.7% correlation improvement in drug sensitivity prediction while achieving comparable performance in gene effect score prediction (Figure 2C). These results were further supported by external validation datasets, where InsilicoCell outperformed all four baseline models with a 13.7% RMSE improvement in drug-protein binding affinity prediction (Figure 2B), as well as outperformed TransCell with a 32.4% correlation improvement in drug sensitivity prediction (Figure 2C). In addition, InsilicoCell outperformed GPS^8^ and DeepCE^7^ in predicting drug-induced gene expression change, as detailed in the following section.

We next assessed the contribution of multi-task learning. Compared with an identical architecture trained separately for each single task, multi-task InsilicoCell achieved superior performance across most tasks, especially with a 24.8% RMSE improvement in drug sensitivity prediction (Figure 2D). A representative example was drug sensitivity prediction in HeLa cell line, where multi-task training elevated the correlation between prediction and ground truth from 0.68 to 0.85 (Figure 2E). Visualization of the last hidden state of drug representations from the transformer encoder blocks revealed that multi-task training generated latent drug embeddings which more faithfully captured the gradient of actual drug sensitivity, while the single-task model failed to recover this structure (Figure 2F). Additionally, we observed performance improvements with increasing training data size, consistent with scaling behavior and suggesting that broader data coverage contributes substantially to model performance (Figure S2).

To dissect the contribution of key model components, we further performed a series of ablation studies. Replacing the transformer module with either a multi-layer perceptron (MLP) or random forest while retaining the same input representations reduced performance across seven tasks (Figure 2G). The largest improvement over MLP was observed for CNV prediction, where InsilicoCell achieved a 18.8% correlation improvement and 49.7% RMSE improvement compared to MLP (Figure 2G). Relative to random forest, InsilicoCell showed even larger performance improvement for all seven tasks. For instance, for drug sensitivity prediction, InsilicoCell improved the average cell line-wise correlation from 0.56 to 0.90 (Figure 2G). Additional ablation analyses showed that current InsilicoCell configuration with task-specific training, dynamic task loss weighting, and the current cell-line and drug representation methods (compared to other SOTA representation methods such as scGPT, GeneFormer, scVI, and GraphFP) performed the best among all tested model configurations (Figure S3, Methods). Under the more stringent entity-level holdout setting, current InsilicoCell also achieved better or comparable prediction performance across all cellular functional profiles compared to other model configurations (Figure S4). Remarkably, InsilicoCell maintained high prediction accuracy for drug sensitivity and gene effect score with average correlation exceeding 0.8 across unseen cell lines (Figure S4, examples in Figure S5).

Taken together, these results demonstrate that InsilicoCell can accurately predict diverse cellular functional profiles and generalizes well to unseen entities. Moreover, the transformer architecture, the two-stage training strategy, dynamic task loss weighting, and modality-specific representations collectively contribute to the overall performance improvement.

### InsilicoCell supports virtual drug screening and drug-mechanism interpretation based on predicted transcriptional responses

A central objective of InsilicoCell is to enable virtual screening of compounds that modulate transcriptional phenotypes across diverse biological contexts. We therefore performed a comprehensive evaluation of its performance in predicting drug-induced transcriptional responses and investigated its utility for downstream applications, including transcriptomics-based drug discovery and mechanism-of-action (MOA) interpretation^1,8,35–37^, before deploying the framework to real-world drug discovery challenges.

We first compared InsilicoCell with representative SOTA models including GPS and DeepCE for predicting drug-induced gene expression changes. An advancement of InsilicoCell is the expansion of predictable cellular contexts, genes, and experimental conditions in comparison to these prior approaches (Figure 3A). InsilicoCell outperformed DeepCE in both sample-level and drug-level holdout settings, achieving performance improvements of 66.1% and 29.2%, respectively (Figure 3B). We further compared InsilicoCell with GPS, a transcriptomics-based drug screening pipeline limited to four cell lines^8^. InsilicoCell achieved superior performance across all evaluated cell lines (Figure 3C, Methods). To further assess model generalizability, we evaluated InsilicoCell on GEOMeta, an external dataset comprising 2,624 drug-induced transcriptional signatures spanning approximately 20,000 genes across diverse biological contexts. Due to differences in data generation and processing, zero-shot prediction yielded only moderate performance; however, supervised fine-tuning substantially improved accuracy (Figure S6A). After observing that prediction performance increased with perturbation magnitude score (Figure S6B, Methods), we chose those signatures with high perturbation magnitude scores to fine-tune InsilicoCell, achieving F1 scores exceeding 0.5 in both signature-level and drug-level holdout settings (Figure S6C, S6D). Because GPS and DeepCE do not explicitly model cellular context, they cannot generalize to the diverse biological contexts represented in GEOMeta.

**Figure 3:**
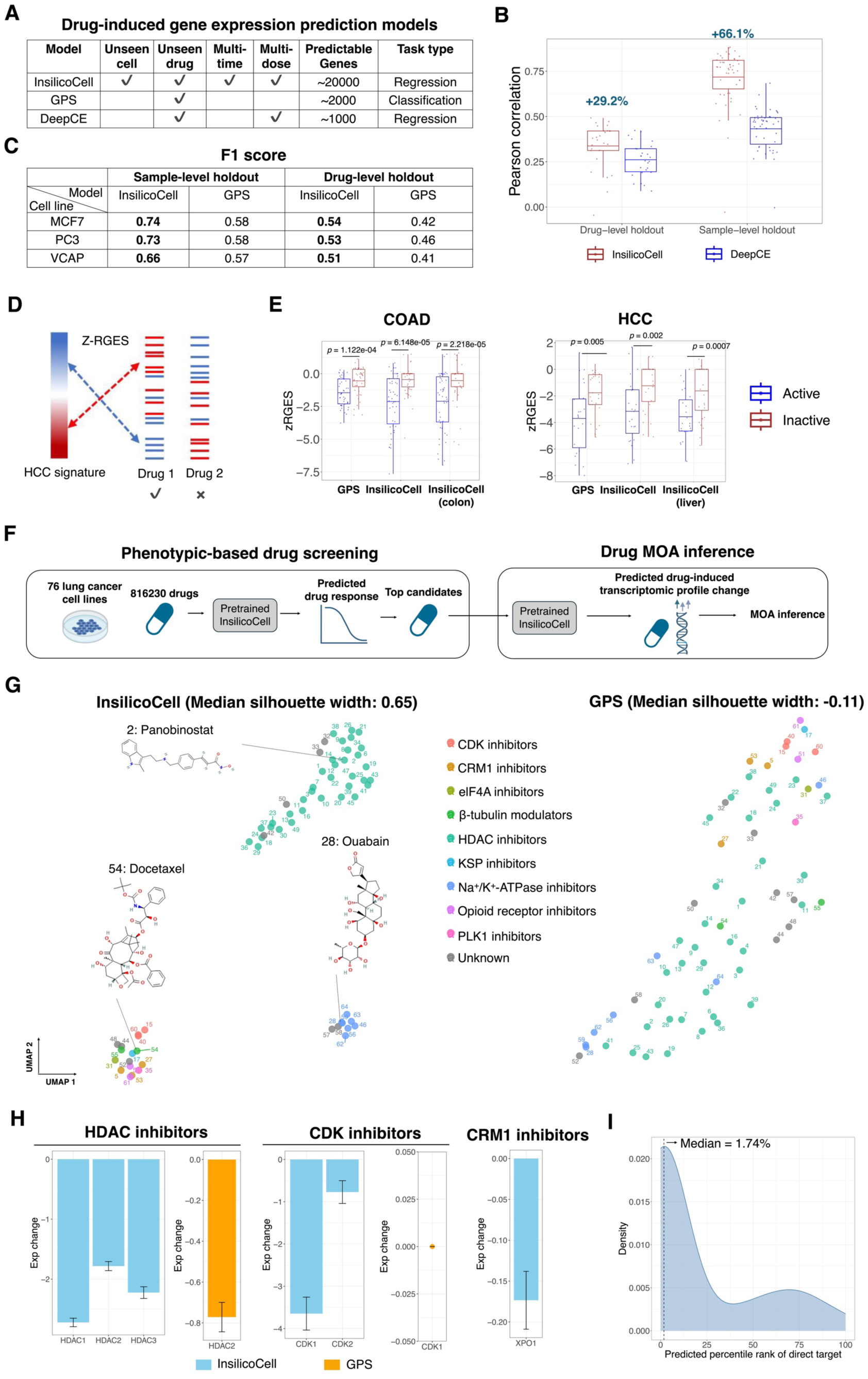
InsilicoCell improves prediction of drug-induced gene expression profiles for virtual drug screening and MOA interpretation. **A**, Comparison of the coverage of predictable genes, cellular contexts, and biological conditions between InsilicoCell and baseline models used for drug-induced gene expression change prediction. **B**, Comparison of the prediction performance on the drug-induced gene expression change task between InsilicoCell and DeepCE. Each dot represents a cell line. Percentage numbers represent changes in mean cell line-wise performance of InsilicoCell compared to DeepCE. **C**, Comparison of the prediction performance on the drug-induced gene expression change task between InsilicoCell and GPS. The continuous prediction values from InsilicoCell were discretized to match the categorical outputs of GPS. F1 score was used for evaluation. **D-E**, Application of InsilicoCell-predicted drug-induced gene expression to virtual drug screening based on disease transcriptomic reversal. **D**, Illustration of the reversal score Z-RGES. Compounds with lower Z-RGES are more effective in reversing disease expression signatures. **E**, Boxplots showing the difference in z-RGES between active and inactive compounds in COAD and HCC. z-RGES was computed from drug-induced gene expression change values predicted by GPS or InsilicoCell using either the same fixed cellular contexts as GPS, or disease-specific cellular contexts (labeled as “colon” or “liver” in the boxplots). Statistical *P* values were computed via Wilcoxon rank-sum tests. **F-I**, Comparison of drug MOA interpretation from the predicted drug-induced gene expression between InsilicoCell and GPS under a virtual drug screening setting. **F**, A total of 816,230 compounds were screened using InsilicoCell-predicted drug sensitivity in lung cancer cell lines. Top candidates with predicted strong drug sensitivity were selected. InsilicoCell or GPS was used to predict drug-induced gene expression change for these selected candidates for further MOA interpretation. **G**, Visualization of drug-induced gene expression change predicted by InsilicoCell and GPS, respectively, for all candidates. Color indicates known MOA classes. **H**, Drug-induced expression change predicted by InsilicoCell or GPS for marker genes in the corresponding MOA group of compounds. **I**, Distribution of rank percentiles for established protein targets among all evaluated proteins by InsilicoCell-predicted drug-protein binding affinity.

Next, we assessed whether InsilicoCell-predicted drug-induced gene expression could support virtual compound screening through the reversal of disease-associated gene expression, as GPS did (Figure 3D, 3E). We compared the screening performance of InsilicoCell to GPS in colon adenocarcinoma (COAD) and HCC. We quantified the reversal using z-transformed reversal gene expression score (z-RGES)^8^, a scoring function used for measuring the extent to which predicted drug-induced gene expression changes oppose a disease-associated transcriptomic signature (Figure 3D). Unlike GPS, which is restricted to four predefined cell lines, InsilicoCell enables drug-induced expression prediction across diverse, disease-relevant cellular contexts. Consistent with the GPS result, z-RGES values derived from InsilicoCell-predicted transcriptional responses significantly distinguished known active from inactive compounds in both COAD and HCC. Moreover, InsilicoCell achieved stronger separation between the two groups than GPS, as reflected by lower P values in both diseases (Figure 3E).

We further explored whether predicted gene expression changes could facilitate MOA interpretation. As a pilot study, we predicted drug sensitivity for 816,230 compounds across 76 lung cancer cell lines, including 51 small cell lung cancer (SCLC) and 25 non-small cell lung cancer (NSCLC) cell lines (Figure 3F, S7). The top 50 compounds predicted to be most potent in NSCLC were subsequently analyzed using their predicted transcriptional responses (Table S2). Compounds sharing the same MOA clustered together in InsilicoCell-predicted transcriptional space, with a median silhouette width of 0.65 outperforming −0.11 from GPS-predicted transcriptional space, indicating that InsilicoCell-predicted transcriptional responses capture biologically meaningful patterns associated with compound activity (Figure 3G). In addition, many of the top candidates identified by InsilicoCell belong to actively-investigated therapeutic classes in NSCLC, such as HDAC inhibitor, CDK inhibitors, and Na+/K+-ATPase inhibitors^38–40^. At the gene level, InsilicoCell recapitulated known target-specific transcriptional effects, including downregulation of *HDAC1 / HDAC2 / HDAC3* by HDAC inhibitors, *XPO1* by CRM1 inhibitors, and *CDK1 / CDK2* by CDK inhibitors (Figure 3H). Several of these genes were either inaccurately predicted or outside the prediction range of GPS. Consistent with these findings, the known targets of most prioritized compounds ranked within the top 2% of all 6,564 proteins in BindingDB based on InsilicoCell-predicted binding affinities (Figure 3I).

Taken together, the results demonstrated the superiority of InsilicoCell compared to other baseline models in predicting drug-induced gene expression change. By extending the range of predictable cellular contexts, genes, and experimental conditions, InsilicoCell enables both transcriptomics-based compound discovery and mechanistic interpretation. Having established its performance through extensive in silico benchmarking, we next applied InsilicoCell to real-world drug discovery challenges.

### InsilicoCell enables multi-objective screening for novel c-Myc activity inhibitors in liver cancer

Conventional target-based drug screening workflows typically begin with biochemical or enzyme-based screening, followed by validation in cellular systems and subsequent confirmation of phenotypic activity. These sequential steps are both time- and resource-intensive and increase the risk of attrition at later stages. By integrating these separate stages into a unified framework, InsilicoCell enables concurrent multi-objective compound screening and early prioritization of candidate compounds, thereby reducing the likelihood of downstream failure. To demonstrate the application of InsilicoCell as a unified virtual platform for multi-objective drug discovery, we tackled three challenging discovery programs. The first case focused on the identification of novel c-Myc activity inhibitors in liver cancer. *MYC* is amplified in many cancers including HCC yet remains one of the most challenging therapeutic targets in oncology^41^. We therefore defined three screening objectives for candidate compounds: (1) inhibition of HCC cell growth, (2) reduction of *MYC* expression in HCC, and (3) inhibition of c-Myc activity.

Before initiating compound screening, we first used InsilicoCell to predict genetic dependency of *MYC* across all HCC cell lines available in DepMap. The results suggested that *MYC* is broadly essential in HCC cell lines (Table S3). Among them, HepG2, one of the most widely used HCC cell lines, exhibited an intermediate level of predicted *MYC* dependency, and was therefore selected as a representative cell line for multi-objective screening. Owing to its multi-modal representation module, InsilicoCell enables scalable screening of large compound libraries such as the Enamine HTS library comprising approximately 1.8 million compounds, the vast majority of which were absent from the pretraining dataset. For every compound, InsilicoCell predicted: (1) drug sensitivity in HepG2 cells, (2) drug-induced *MYC* expression change in HepG2 cells, and (3) inhibition effects on c-Myc activity (Figure 4A).

**Figure 4:**
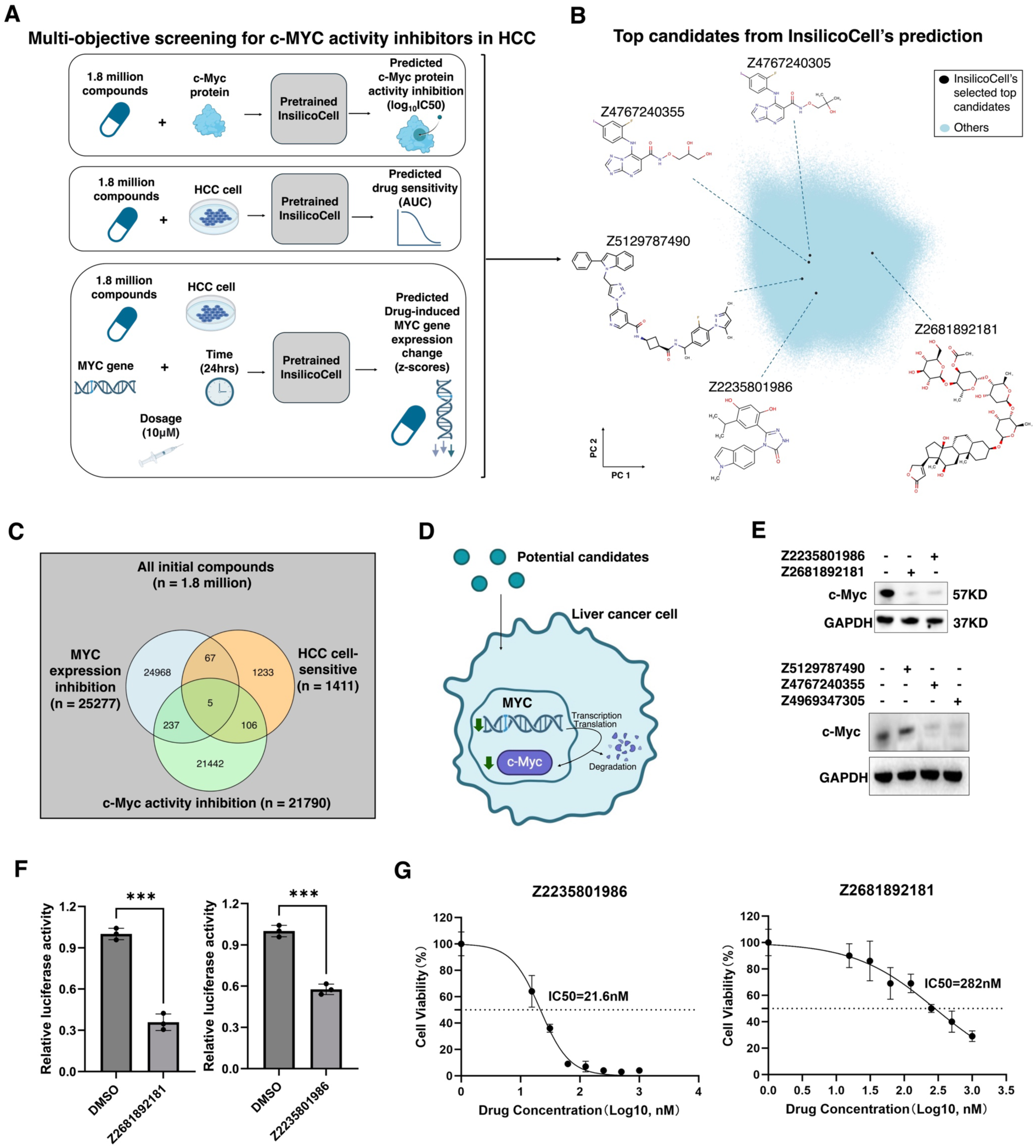
InsilicoCell enables multi-objective screening for discovering new c-Myc activity inhibitors in HCC. **A**, Overview of applying InsilicoCell to multi-objective screening for c-Myc activity inhibitors in HCC. Three objectives were included: (1) inhibiting c-Myc protein activity, (2) inhibiting cell growth in HCC, and (3) downregulating *MYC* expression in HCC. **B**, Chemical structure space of library compounds, highlighting the final selected candidates: Z5129787490, Z4767240355, Z4767240305, Z2235801986, Z2681892181. **C**, Number of compounds predicted to satisfy each screen objective, out of all 1.8 million screened compounds. **D**, Schematic of the major biological processes represented by the three objectives. Upon entering the liver cancer cell, candidate compounds were predicted to perturb MYC-related processes, including transcription, translation and protein degradation. **E**, Western blot showing c-Myc protein expression reduction in HepG2 cells under treatment of the top drug candidates selected by InsilicoCell. **F**, Dual-luciferase reporter assay showing c-Myc protein activity inhibition with Z2235801986 and Z2681892181 (t-test: *: < 0.05, ** < 0.01, *** < 0.001). **G**, Dose-response curves of Z2235801986 and Z2681892181 in HepG2 cells.

Among the 1.8 million screened compounds, thousands satisfied at least one of the three screening criteria, and several hundred satisfied two screening criteria. Ultimately, only five compounds satisfied all three criteria (Figure 4B-C, S9A). Compared to the conventional sequential screening strategies which focus on only a single objective at a time and often advance compounds that subsequently fail at later stages, InsilicoCell enabled coordinated multi-objective prioritization by jointly evaluating all screening criteria, thus largely reducing the compound failure rate (Figure 4C). By meeting all three objectives, the ultimately selected candidates were expected to affect the biological processes of *MYC* transcription, translation and c-Myc degradation upon entering the liver cancer cell (Figure 4D). Further, the structures of the five candidate compounds are not similar to known c-Myc activity inhibitors from our pretrained data (Figure S8), indicating the novelty of the predicted compounds.

To validate the candidates selected by InsilicoCell, we conducted in vitro experiments in HepG2. Western blot results confirmed that all five compounds reduced c-Myc levels, with four compounds (Z4767240355, Z4969347305, Z2235801986 and Z2681892181) nearly abolishing c-Myc expression (Figure 4E). Consistent with these findings, dual-luciferase reporter assay results showed that Z2235801986 and Z2681892181 significantly inhibited c-Myc protein transcriptional activity (Figure 4F), while the other three compounds also showed activity inhibition effects at higher concentrations (Figure S9B). All five compounds inhibited cell growth in HepG2 cells (Figure 4G, S9C), where Z2235801986 and Z2681892181 showed particularly high potency, with IC_50_ of 21.6 nM and 282 nM, respectively (Figure 4G). The other two compounds, Z4767240355 and Z4969347305 showed dose responses while exhibited weaker short-term growth inhibition (Figure S9C). Further cell colony formation assays revealed significant suppression of long-term HepG2 cell growth following treatment at 25 *μ*M (Figure S9D-S9F). Taken together, these results validate the predictions of InsilicoCell and demonstrate the feasibility of its multi-objective screening approach for the de novo discovery of c-Myc activity inhibitors in HCC.

### InsilicoCell enables multi-objective screening for novel anti-fibrotic inhibitors in idiopathic pulmonary fibrosis

To demonstrate the application of InsilicoCell to more diverse disease and cellular contexts, we applied InsilicoCell to the discovery of compounds targeting myofibroblasts in IPF based on the single-cell transcriptomics^42^ (Figure 5A). IPF is a chronic lung disease arising from lung injury and the subsequent fibrotic response, leading to thickened alveolar walls and the loss of normal alveolar space, while the etiology and the role of microenvironments in its disease progression remain largely unknown^43^. Myofibroblasts are the ultimate effector cells driving fibrotic tissue modeling and their accumulation is a hallmark of IPF. These cells originate majorly from resident fibroblasts whose phenotypic differentiation is driven by profibrotic mediators such as TGF-*β* (Figure 5B). Reduced expression of canonical profibrotic markers, including ACTA2 (α-SMA), COL1A1, and FN1, is widely regarded as an indicator of myofibroblast deactivation and phenotypic reversion toward a more quiescent fibroblast state. This transition is associated with attenuation of extracellular matrix deposition and restoration of tissue homeostasis, thereby contributing to the resolution or reversal of fibrotic remodeling in the lung^43–45^.

**Figure 5:**
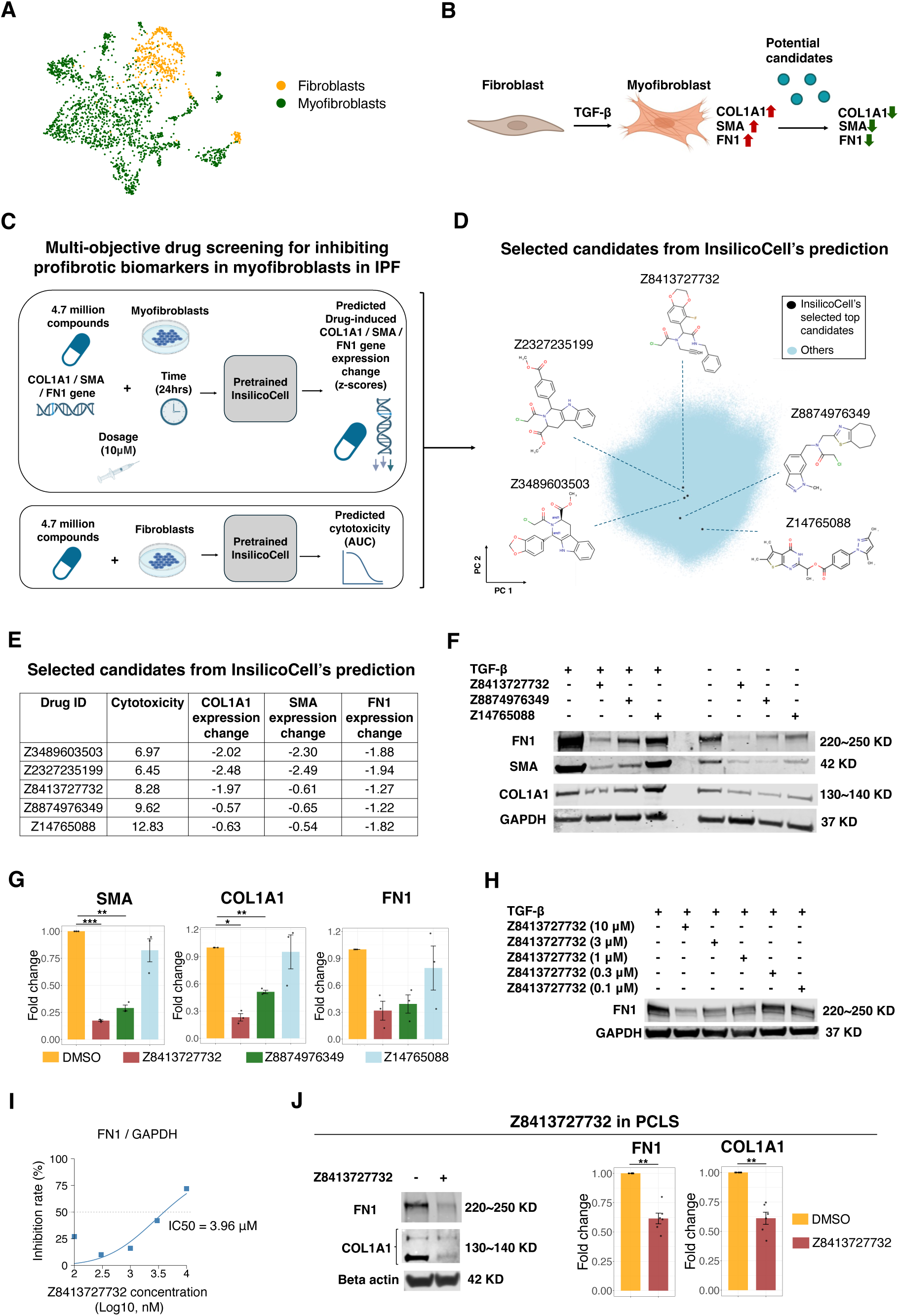
InsilicoCell enables multi-objective screening for profibrotic marker suppression in IPF. **A**, Visualization of myofibroblasts and fibroblasts in a single-cell transcriptomic dataset from IPF. **B**, Schematic of IPF in vitro model. Myofibroblasts are induced from fibroblasts via TGF-*β* treatment. Potential drug candidates are expected to downregulate three key fibrosis marker genes in myofibroblasts. **C**, Overview of applying InsilicoCell to multi-objective screening for profibrotic marker suppressors in myofibroblasts. Two objectives were included: (1) downregulating expression of COL1A1, SMA, and FN1 in myofibroblasts, and (2) low cytotoxicity in fibroblasts. **D**, Chemical structure space of the compound library, highlighting the ultimately selected candidates: Z3489603503, Z2327235199, Z8413727732, Z8874976349, Z14765088. **E**, InsilicoCell-predicted values based on the two objectives for the five selected candidates. Predicted drug-induced gene expression change values were reported as z-scores and predicted cytotoxicity values were represented by AUC values. **F**, Western blot showing reduced expression of fibrosis markers COL1A1, SMA, and FN1 in TGF-*β* induced myofibroblasts under treatment of Z8413727732, Z8874976349, and Z14765088. **G**, Boxplots showing three independent western-blot experiments in myofibroblasts. Ratio paired t-test was used for statistically measuring the difference in the expression level of fibrosis markers between the compound candidates and the DMSO control (*: < 0.05, ** < 0.01, *** < 0.001). **H-I**, Extended exploration of dose-dependent effect of Z8413727732. **H**, Western blot showing reduced expression of fibrosis markers in myofibroblasts under treatment of Z8413727732 at various dosages. **I**, Dose-response curve showing inhibition effect of Z8413727732 in myofibroblasts. **J**, Extended exploration of inhibition effect on FN1 and COL1A1 by Z8413727732 in PCLS. Boxplots showed western-blot experiments on six independent samples of PCLS from human IPF lung tissue under 10μM Z8413727732 treatment.

Because currently approved therapies provide only limited efficacy and do not adequately reverse established fibrosis, we sought to identify novel anti-fibrotic compounds capable of suppressing multiple profibrotic markers while minimizing toxicity to fibroblasts. Specifically, we defined a multi-objective screening strategy that prioritized compounds predicted to (1) reduce the expression of COL1A1, ACTA2 (α-SMA), and FN1 in myofibroblasts while (2) exhibiting low cytotoxicity (represented as drug sensitivity) in fibroblasts. To achieve this, we virtually screened 4.7 million compounds from the combined Enamine libraries (Methods). For each compound, InsilicoCell predicted the following: (1) drug-induced *COL1A1 / SMA / FN1* gene expression change in myofibroblasts, and (2) drug sensitivity in fibroblasts (Figure 5C). Ultimately, five candidates were selected for bench validation (Figure 5D), and their predicted values for the two objectives are summarized in Figure 5E. Among them, the two compounds Z3489603503 and Z2327235199, despite predicted downregulation on the three fibrosis markers, were predicted to be highly cytotoxic to fibroblasts (Figure 5E). The other three compounds were predicted to be less cytotoxic to fibroblasts, making them more favorable candidates for further evaluation (Figure 5E).

These five candidates were then acquired for validation using the standard in vitro fibrosis model, where myofibroblasts were induced from fibroblasts by TGF-*β* treatment (Figure 5B). Consistent with model prediction, Z3489603503 and Z2327235199 were remarkedly cytotoxic in fibroblasts, as observed by microscopy, precluding further assessment of fibrosis marker expression due to extensive cell death in both myofibroblasts and fibroblasts at 10*μ*M (Figure S10). For the other three candidates, which were not cytotoxic in fibroblasts, western blot results showed that Z8413727732 and Z8874976349 significantly reduced fibrotic marker expression in myofibroblasts (Figure 5F, 5G). Z14765088 also showed some inhibition effects, although to a lesser extent.

To further investigate the anti-fibrotic effects of the most effective compound, Z8413727732, we conducted extended study to quantify the dose-dependent inhibition effect of Z8413727732 in myofibroblasts (Figure 5H, 5I). Z8413727732 suppressed the marker expression in a dose-dependent manner, with an estimated IC_50_ of 3.96 *μ*M (Figure 5I). Notably, compared to two FDA-approved IPF drugs, Z8413727732 showed more effective inhibition on the three fibrotic markers than nerandomilast, and similarly effective as nintedanib at the same dose of 10*μ*M (Figure S11A). Furthermore, to evaluate the anti-fibrotic effects of Z8413727732 in a more clinically relevant setting, we extended our studies from in vitro cell cultures to ex vivo precision-cut lung slices (PCLS) from six independent samples collected from human IPF lung tissues. Western-blot results confirmed the inhibition effects of Z8413727732 on FN1 and COL1A1 expression ex vivo (Figure 5J). Mechanistically, Z8413727732 inhibited phosphorylation of SMAD3, a key downstream mediator of TGF-*β* and a transcriptional activator of the fibrotic effectors COL1A1, SMA and FN1, suggesting that Z8413727732 may suppresses myofibroblast activation, at least in part, through the TGF-*β*/SMAD3 signaling axis (Figure S11B). Taken together, our results validate the novel discovery made by InsilicoCell on compound candidates suppressing profibrotic markers in myofibroblasts in IPF via its multi-objective drug screening approach. Moreover, this case also demonstrates the feasibility of applying InsilicoCell to single-cell transcriptomic data for prediction.

### InsilicoCell enables multi-objective screening of compounds for inducing pluripotency marker expression in mesenchymal stem cells

In addition to diseased contexts, we next sought to evaluate the multi-objective compound screening capability of InsilicoCell in a healthy cellular context. As the third case study, we investigated the discovery of novel compounds that induce the expression of pluripotency markers in mesenchymal stem cells (MSCs). Generating and maintaining pluripotent cells in vitro has long been a biological and technical challenge. Seminal studies demonstrated overexpression of transcriptional factors OCT4, SOX2, NANOG, and c-Myc to be sufficient for reprogramming human somatic cells into induced pluripotent stem cells (iPSCs)^46,47^. However, conventional iPSC reprogramming strategies rely on inserting exogenous DNA and frequently involve overexpression of the oncogene c-Myc, raising safety concerns for clinical applications^48^. Consequently, the use of small molecules to modulate stem cell potency and replace genetic reprogramming factors has attracted substantial interest^49^. MSCs, despite their low pluripotency level compared to human embryonic stem cells (hESCs), are readily accessible and serve as ethically tractable and clinically relevant alternatives to iPSCs and hESCs^50^. With observed low endogenous expression of key pluripotency regulators *OCT4*, *SOX2*, and *NANOG* in MSCs, identifying compounds that induce their expression represents a promising strategy for enhancing pluripotency potential. However, performing large-scale compound screening in vitro remains technically challenging and resource intensive^51^.

Conventional structure-based drug screening approaches typically optimize compounds against a single target and are not readily suited for simultaneously modulating multiple pluripotency regulators. We therefore formulated a multi-objective screening strategy in MSCs that prioritized compounds predicted to (1) upregulate pluripotency marker expression (*OCT4*, *SOX2*, and *NANOG*) while (2) maintaining low cytotoxicity in MSCs. We used the same compound library as the IPF case study. For each compound, InsilicoCell predicted the following: (1) drug-induced *OCT4* / *SOX2* / *NANOG* gene expression change in MSCs, and (2) drug sensitivity in MSCs (Figure 6A). As a result, five candidates were selected for bench validation (Figure 6B), and their predicted values for the two objectives are shown in Figure 6C. The five compounds were experimentally evaluated in Wharton’s jelly-derived mesenchymal stem cells (WJMSCs) (Methods). Consistent with model prediction, these compounds generally supported healthy cell growth with no obvious cell death (Figure 6D). Among them, Z1903071952 and Z2964511831 were validated by qPCR experiments to significantly induce the pluripotency markers in WJMSCs after 48-hour treatment under 10*μ*M (Figure 6E). Although chemical reprogramming of cell states typically requires prolonged treatment, stage-specific interventions, and combinations of multiple compounds^52^, our pilot study demonstrates the feasibility of applying InsilicoCell to identify promising initial candidates for such strategies, which would be challenging to obtain through conventional experimental screening approaches.

**Figure 6:**
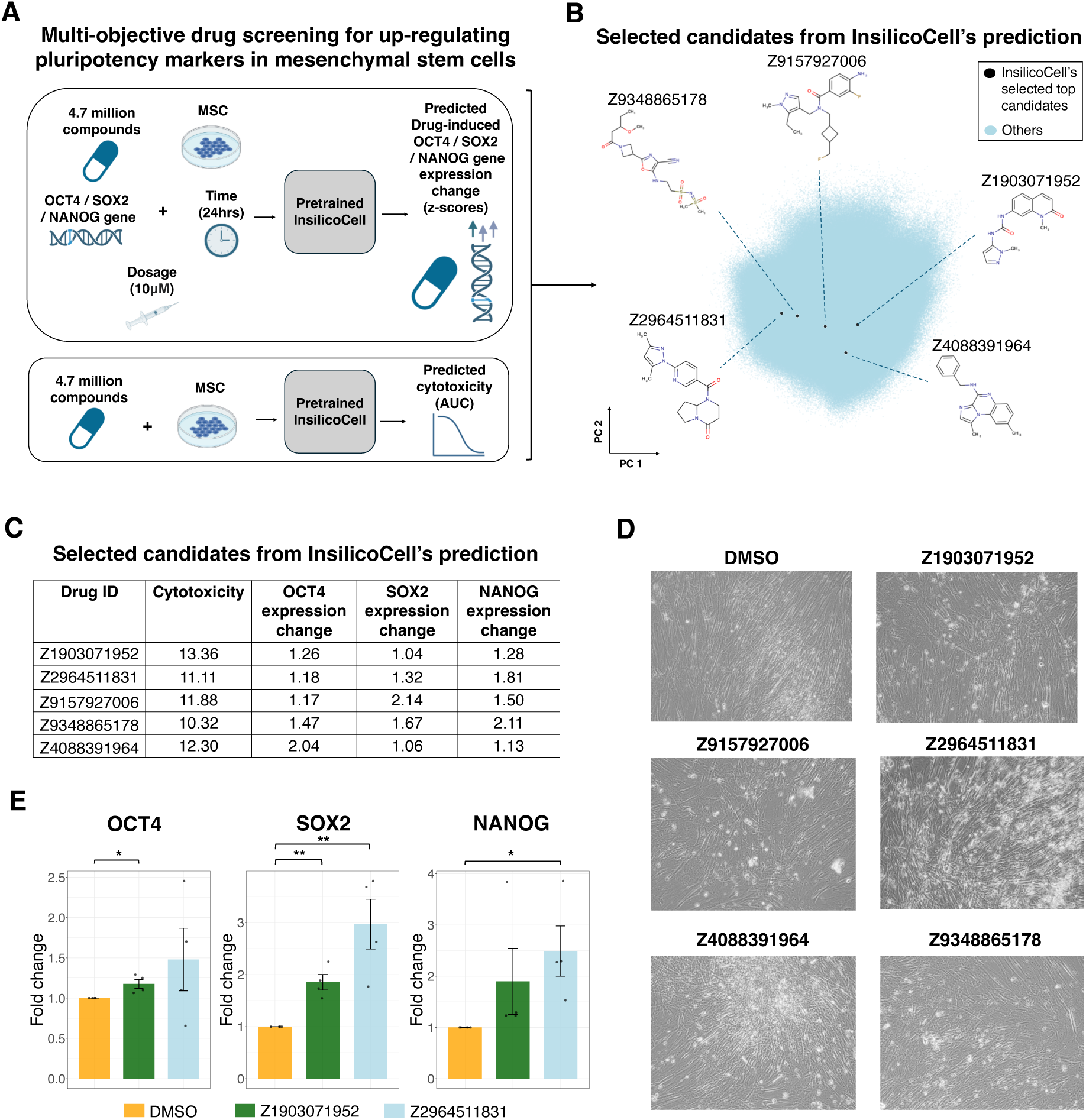
InsilicoCell enables multi-objective screening for inducing pluripotency markers in mesenchymal stem cells. **A**, Overview of applying InsilicoCell to multi-objective screening for pluripotency marker inducers in MSCs. Two objectives were included: (1) upregulating expression of *OCT4*, *SOX2*, and *NANOG* in MSCs, and (2) low cytotoxicity in MSCs. **B**, Chemical structure space of the compound library, highlighting the ultimately selected candidates: Z1903071952, Z2964511831, Z9157927006, Z9348865178, Z4088391964. **C**, InsilicoCell-predicted values based on the two objectives for the five selected candidates. Predicted drug-induced gene expression change values were reported as z-scores and predicted cytotoxicity values were represented by AUC values. **D**, Microscopy images showing survival of MSCs treated with 10*μ*M of Z1903071952, Z2964511831, Z9157927006, Z9348865178, and Z4088391964 for 48 hours. **E**, Boxplots showing four independent qPCR experiments in WJMSCs treated with Z1903071952 or Z2964511831. Ratio paired t-test is used for statistically measuring the difference in the expression level of pluripotency markers between the candidate compounds and the DMSO control (*: < 0.05, ** < 0.01, *** < 0.001).

### InsilicoCell supports patient-level and spatial-level prediction and new task addition through domain adaptation

Beyond predicting cellular functional profiles for cell lines and single cells (Figure 4-6), we further investigated the potential of extending InsilicoCell to patient-level and spatially resolved applications in translational research. A significant application is the prediction of patient-specific drug sensitivity. Unlike cell lines, rich patient-level drug sensitivity data are difficult to collect, making it challenging to train a model directly in patient cohorts. Directly using a model trained on cell lines to make zero-shot prediction on patients may experience domain shift, leading to reduced performance. To address this limitation, we investigated a few-shot learning framework, in which InsilicoCell leverages a small number of experimentally annotated samples from the target domain to improve performance. Starting with patient data without any sensitivity data, the framework employs active learning to identify the most informative drug-patient pairs for experimental measurement. The resulting labels are then used to fine-tune the pretrained model, enabling efficient adaptation to the new domain while minimizing experimental costs^53^.

To demonstrate zero-shot prediction and few-shot adaptation at the patient level (Figure 7A), we evaluated InsilicoCell on the BeatAML dataset, which contains pretreatment blood transcriptomes from patients with acute myeloid leukemia (AML) and matched ex vivo drug sensitivity measurements^54^ (Table S1, Methods). Across all evaluated fine-tuning set sizes (300 to 1000 drug-patient pairs), the active learning-based few-shot approach consistently outperformed the random sampling-based few-shot approach on the test set (Figure 7B). When using 1000 selected drug-patient pairs for fine-tuning, compared to the random sampling-based approach, the active learning-based approach improved the averaged patient-wise correlation and RMSE by 17.1% and 11.1%, respectively (Figure 7B); compared to zero-shot prediction, the improvements were even more pronounced, with gains of 45.4% and 39.1%, respectively, in the averaged patient-wise correlation and RMSE (Figure 7B). These results demonstrate that InsilicoCell can be effectively transferred to patient-level applications through zero-shot prediction and that its active learning-guided few-shot adaptation framework substantially improves performance in data-limited clinical settings.

**Figure 7:**
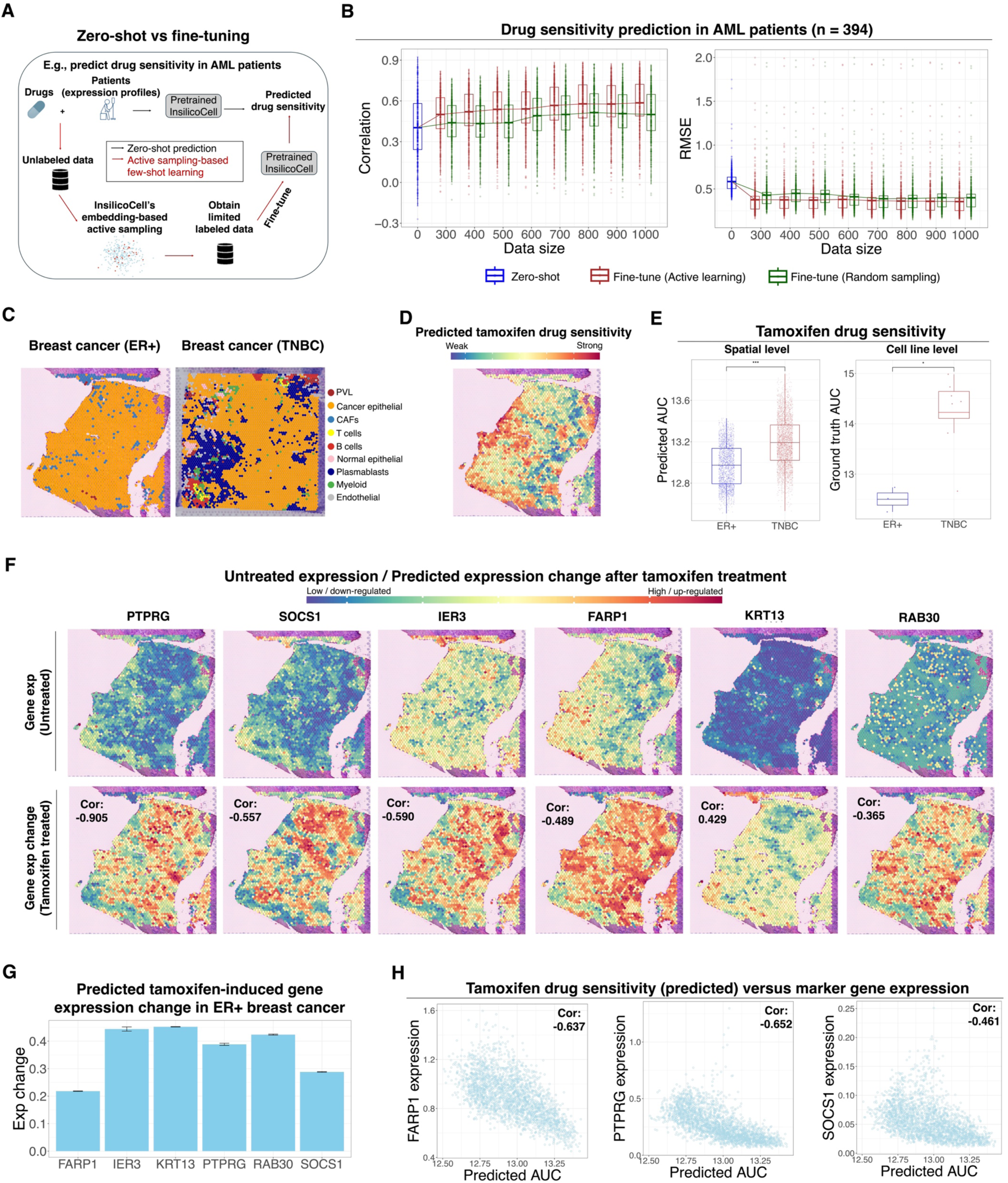
Application of InsilicoCell to patient-level and spatial-level prediction via domain adaptation. **A-B**, An example usage of InsilicoCell for predicting drug sensitivity in AML patients. **A**, Both zero-shot prediction and few-shot adaptation are applicable. For few-shot adaptation, InsilicoCell adopts an active learning-based sampling method for selecting the optimal samples of drug-patient pairs for drug sensitivity label acquisition and fine-tuning the model. **B**, Comparison of prediction performance on the test set between zero-shot prediction and few-shot adaptation (fine-tuning). Sampling methods were compared between active learning-based sampling and random sampling, under the same number of selected samples for fine-tuning. Each dot represents a patient (n = 394). **C-H**, An example usage of InsilicoCell for predicting drug sensitivity and drug-induced gene expression change for tamoxifen based on spatial transcriptomes of breast cancer. **C**, Visualization of cell types and their locations on the tissue samples. **D**, Visualization of tamoxifen drug sensitivity values (AUC) predicted by InsilicoCell on the ER+ breast cancer tissue. **E**, Boxplots showing the spatial-level drug sensitivity values predicted by InsilicoCell (left), and the cell line-level ground truth of drug sensitivity from CTRP database (right). Each dot represents a location on the tissue (left) or a cell line (right) (Wilcoxon rank-sum test: *: < 0.05, ** < 0.01, *** < 0.001). **F**, Visualization of the original untreated expression and the tamoxifen-induced gene expression change (z-scores) predicted by InsilicoCell for six reported gene markers robustly induced by tamoxifen in ER+ breast cancer. **G**, Bar plots showing the InsilicoCell-predicted average expression change values across all locations on the ER+ breast cancer tissue after tamoxifen treatment for the six genes. **H**, Scatter plots showing the correlation between the original untreated expression for three gene markers and the tamoxifen drug sensitivity across all locations as predicted by InsilicoCell.

Beyond patient-level prediction, we investigated whether InsilicoCell could be extended to spatial transcriptomics (ST). Unlike bulk or single-cell RNA-seq, ST preserve spatial context, enabling the study of spatially organized transcriptomic landscape in situ^55^. However, spatially resolved functional profiling remains technically challenging. InsilicoCell addresses this gap by enabling prediction of in situ cellular functional profiles, including drug sensitivity and drug-induced transcriptional responses, directly from ST data through unsupervised domain adaptation (Methods).

Because spatially resolved functional measurements are not yet widely available, we evaluated the spatial prediction capability of InsilicoCell using tamoxifen response in breast cancer as a case study (Figure 7C-7H). Tamoxifen is a selective estrogen receptor (ER) modulator used for ER+ breast cancer treatment^56^. The 10x Visium ST data of ER+ breast cancer and triple-negative breast cancer (TNBC) tissue samples were obtained from a published study^57^. Cell type annotation revealed that most spatial spots were dominated by malignant epithelial cells (Figure 7C). InsilicoCell was first used to predict per-spot tamoxifen drug sensitivity from the ST data for both ER+ breast cancer and TNBC tissue samples (Figure 7D, 7E left). Cancer cells in ER+ breast cancer were predicted to be significantly more sensitive to tamoxifen than those within TNBC tumors (Figure 7E, left), consistent with the established mechanism of action of tamoxifen and with experimentally measured differences in drug sensitivity between ER+ and TNBC cell lines (Figure 7E, right). These results indicate that InsilicoCell captures biologically meaningful, subtype-specific drug sensitivity patterns in a spatial context.

We further utilized InsilicoCell to predict per-spot tamoxifen-induced gene expression change (z-scores) in ER+ breast cancer tissue. We focused on six genes (*PTPRG*, *SOCS1*, *IER3*, *FARP1*, *KRT13*, and *RAB30*) previously reported to be robustly upregulated by tamoxifen treatment^56^. Across all spatial spots, predicted tamoxifen-induced expression changes highly correlated with baseline (untreated) expression levels for all six genes (Figure 7F), indicating that InsilicoCell sensitively captured the spatial expression variation across cells during prediction. The average expression change values among all spots showed that all six genes were upregulated by tamoxifen in the ER+ breast cancer tissue (Figure 7G), which aligns with the reported study^56^. These results indicate that InsilicoCell can recapitulate known tamoxifen-responsive transcriptional programs while preserving spatial variation in predicted drug responses across tissue regions.

Interestingly, we also found InsilicoCell-predicted tamoxifen drug sensitivity across all spots (Figure 7D) was negatively correlated with the baseline expression of several genes (*FARP1*, *PTPRG*, *SOCS1* as in Figure 7F) (Figure 7H). This suggests the potential of *FARP1*, *PTPRG*, and *SOCS1* as marker genes for inferring tamoxifen sensitivity in ER+ breast cancer, which has not been reported. More broadly, it suggests that InsilicoCell-predicted in situ tamoxifen drug sensitivity is likely to capture the underlying gene regulatory and signaling mechanisms. Taken together, our results demonstrate the potential of InsilicoCell for patient-level and spatial-level cellular functional profiles prediction via supervised and unsupervised domain adaptation (Figure 7A-7H).

In addition to the seven types of cellular functional profiles included in InsilicoCell, we explored the potential of adapting the pretrained InsilicoCell to new tasks via transfer learning (S12A, Methods). As a representative example, we considered drug combination synergy, a new type of cellular functional profiles not included in InsilicoCell during pretraining. Our results suggest that the representations learned by InsilicoCell are sufficiently generalizable to support adaptation to related cellular prediction tasks beyond those explicitly included during pretraining (Figure S12B, Methods).

## Discussion

Recent advances in foundation models across disciplines, coupled with the rapid accumulation of large-scale cellular datasets, have accelerated efforts toward building “virtual cell” models^19,20^. Existing efforts largely fall into two categories: mechanistic models that simulate physical and molecular interactions, typically in simplified cellular systems^21,22,24^, and data-driven models that leverage large-scale single-cell transcriptomics for predicting transcriptional responses to perturbations^58–63^. Realizing the full potential of virtual cell models will require sustained, community-wide collaboration across disciplines to generate data and build models. In contrast, InsilicoCell is motivated by practical applications in experimental biology and drug discovery through leveraging existing diverse, large-scale datasets. A central design principle of InsilicoCell is the ability to generalize across unseen entities, contexts and conditions, support the addition of new tasks and datasets, and meanwhile support the prediction of a wide range of cellular functional profiles within a unified framework.

Guided by the key principle, we introduced several methodological innovations. First, one major limitation of existing approaches is the lack of unified pretrained models capable of predicting multiple types of cellular functional profiles within a single framework. Most models are developed for individual tasks, restricting their ability to generalize and share information across biological layers^4,5,7–15^. Inspired by multi-modal foundation models in other domains^16–18^, InsilicoCell tokenizes diverse biological modalities into a unified representation space, enabling flexible integration of heterogeneous inputs.

Second, modality-specific embeddings allow generalization to previously unseen entities. Examples include chemical fingerprints for compounds, ontology-based representations for genes, and sequence-based representations for proteins. In addition, inspired by previous studies showing transcriptomic data alone can capture substantial information about genomic features and cellular responses^4^, we adopted transcriptomic profiles as a scalable and broadly applicable representation of cellular context. This design enables deployment across diverse biological systems while avoiding reliance on specialized data modalities that remain less widely available. Although integrating additional omics layers could further enhance context representation, their limited availability currently constrains widespread adoption.

Lastly, unlike unsupervised foundation models, InsilicoCell leverages large-scale labeled data for supervised pretraining. Its multi-task transformer architecture enables joint learning across diverse tasks, allowing knowledge transfer and improving performance through shared representations. For example, tasks with limited compounds, such as drug sensitivity prediction, benefit from information learned from related tasks with a substantially larger collection of compounds, such as drug-protein binding prediction, although they are not context-dependent. This shared learning framework enhances both performance and generalization. Furthermore, we observed scaling behavior in InsilicoCell, with performance improving as additional data are incorporated, suggesting that continued expansion of training data will further strengthen its predictive capability.

Moving beyond in silico benchmarking, we applied InsilicoCell to three real-world applications spanning both cancers and non-cancerous diseases. Conventional target-based screening approaches, including structure-based docking, face substantial challenges when applied to proteins considered difficult to target because of the absence of well-defined binding pockets. Identifying compounds that simultaneously modulate multiple targets presents an even greater challenge within these frameworks. In contrast, by extending the transcriptomics-based screening paradigm introduced in GPS, InsilicoCell enables the screening of novel compounds that indirectly up/downregulate the expression of phenotype-associated genes, thereby drive desired phenotypic changes. Furthermore, the unified model predicts additional properties like cytotoxicity, facilitating multi-objective compound prioritization. Experimental validation from three bench labs confirmed the predicted activities of the majority of tested compounds selected from large-scale virtual screens, demonstrating the great potential of applying InsilicoCell to a broad range of disease and cellular contexts, including cancer, non-cancerous diseases and cellular engineering applications.

This study has several limitations. First, InsilicoCell does not currently incorporate lead optimization, which would enable iterative refinement of promising hits to improve their properties. Future work could integrate InsilicoCell with optimization frameworks, such as MolSearch^64^, to support end-to-end drug discovery. Because the identified hits had not undergone medicinal chemistry optimization or selectivity evaluation, we did not pursue extensive preclinical validation studies. Second, the model would benefit from broader and more diverse biological contexts, particularly perturbation data from primary and immune cells and context-specific molecular interactions, to further enhance generalization across cellular systems. As additional labeled datasets become available, InsilicoCell can be expanded to include more modalities, thereby improving predictive performance. Moreover, incorporating additional types of cellular functional profiles, such as proteomic/metabolite/morphological measurements in molecular states, protein-protein interactions in molecular interactions, and genetic-perturbed transcriptional changes in perturbation responses, may further extend the model’s capability. Additionally, although InsilicoCell is currently pretrained on cell lines but not single-cell or patient data due to the scarcity of labeled cellular function profile data in these domains, it is possible to incorporate these data in the future when they become abundant. Finally, InsilicoCell does not explicitly address model interpretability or predictive uncertainty. Although its performance is supported by multiple experimental validations, improving interpretability could provide mechanistic insights into underlying biological relationships, while uncertainty estimation would enable more reliable prioritization of predictions and better account for experimental variability.

## Methods

### Computational Methods

#### Datasets

We collected datasets covering a variety of tasks relating to cellular functional profiles for model training and validation. Details of the datasets are presented in Table S1.

#### Cellular functional profiles

Each of the following cellular functions profiles is viewed as a prediction task:

##### (1) Drug-induced gene expression change

Labeled data were obtained from LINCS (level 5 data)^1^. The expression change values were preprocessed as z-scores in LINCS, where more positive values represent greater upregulation, while more negative values represent greater downregulation of gene expression, relative to control treatments. We selected high-quality signatures with commonly used treatment conditions (time points: 3h, 4h, 6h, 24h, 48h; and dosages: 0.1uM, 1uM, 2.5uM, 5uM, 10uM, 20uM). Each label represents the expression change of a gene (*G*) induced by a drug (*D*) at a time point (*T*) and a dosage (*S*) in a cell line (*C*). Given a total of *N* labels in the dataset, there are a total of *N* samples and the input of the *i*th sample can be represented as [*D_i_, G_i_, C_i_, T_i_, S_i_*] (i ∈ {1, 2, 3,…, N}).

##### (2) Ligand-target binding affinity

Labeled data were obtained from BindingDB^29^. The binding affinity values were reported as IC_50_ in nM, where smaller values represent stronger ligand-protein interactions. We log-transformed the values (log_10_IC_50_) before model training. Each label represents the binding affinity between a drug (*D*) and a protein (*P*). Given a total of *N* labels in the dataset, there are a total of *N* samples and the input of the *i*th sample can be represented as [*D_i_, P_i_*] (i ∈ {1, 2, 3,…, N}).

##### (3) Drug sensitivity

Labeled data were obtained from CTRP using the PharmacoGx package^31,65^. The sensitivity values were reported as area under the dose-response curve (AUC), where lower AUC values indicate stronger growth inhibition effect of a drug in a cell line. We log-transformed the values (ln(AUC+1)) before model training. Each label represents the sensitivity of a cell line (*C*) to a drug (*D*). Given a total of *N* labels in the dataset, there are a total of *N* samples and the input of the *i*th sample can be represented as [*D_i_, C_i_*] (i ∈ {1, 2, 3,…, N}).

##### (4) Gene effect score

We used the preprocessed labeled DepMap data (release version: 19Q3) from the TransCell project^4^. Gene effect scores quantify the effect of gene perturbation on cancer cell viability. More negative scores indicate stronger gene dependency for cell growth or survival. Scores of less than −1 represent strong cell inhibition effects upon gene loss. Each label represents the gene effect score for a knockout/silenced gene (*G*) in a cell line (*C*). Given a total of *N* labels in the dataset, there are a total of *N* samples and the input of the *i*th sample can be represented as [*G_i_, C_i_*] (i ∈ {1, 2, 3,…, N}).

##### (5) TF-target gene association

Labeled data were obtained from ChEA^30^. Each label represents the association between a transcription factor (*F*) and a gene (*G*). The labels are binary where 1 represents a reported TF-gene regulatory association and 0 represents no known association. Given a total of *N* labels in the dataset, there are a total of *N* samples and the input of the *i*th sample can be represented as [*G_i_, F_i_*] (i ∈ {1, 2, 3,…, N}).

##### (6) Gene mutation

We used the preprocessed labeled DepMap data (release version: 19Q3) from the TransCell project^4^. Each label represents the mutation status of a gene (*G*) in a cell line (*C*). The labels are binary where 1 represents the presence of a mutation and 0 represents no detected mutation. Given a total of *N* labels in the dataset, there are a total of *N* samples and the input of the *i*th sample can be represented as [*G_i_, C_i_*] (i ∈ {1, 2, 3,…, N}).

##### (7) CNV

We used the preprocessed labeled DepMap data (release version: 19Q3) from the TransCell project^4^. CNV values represent gene-level copy number relative to a reference baseline. Values near zero correspond to a normal copy number state, whereas positive and negative values indicate copy number gains (amplifications) and losses (deletions), respectively. Each label represents the gene-level (*G*) copy number variation in a cell line (*C*). Given a total of *N* labels in the dataset, there are a total of *N* samples and the input of the *i*th sample can be represented as [*G_i_, C_i_*] (i ∈ {1, 2, 3,…, N}).

#### Input embeddings

##### (1) Drug representation

###### ECFP representation

ECFP4 (1024-bits) features were generated using the rdkit package (https://doi.org/10.5281/zenodo.591637) to represent each drug.

###### GNN representation

We adopted GraphFP^66^ for molecule representation. This method encodes each molecule from two complementary views: the molecular graph and the fragment graph, where fragments represent substructures derived from the parent molecule. Both graphs were independently processed using separate GNNs. To align the two views, a contrastive objective encourages the fragment embedding from the fragment GNN to match the aggregated embedding of its corresponding atoms from the molecular GNN. The final molecule embedding was obtained by aggregating all fragment-level embeddings from both views. Following the original GraphFP design, we used a 5-layer GIN^67^ for molecular graphs and a 2-layer GIN for fragment graphs, both with a hidden dimension of 300 and mean pooling for aggregation. The model was pretrained on the training split of all combined molecules after removing invalid structures, resulting in 704,943 molecules. Pretraining was conducted for 300 epochs using the AdamW optimizer (initial learning rate 2×10^-3^, batch size 2048), with the learning rate decayed by 0.8 every 5 epochs without improvement. All pretraining was performed on a single NVIDIA RTX A5000 GPU and required approximately 15 hours.

##### (2) Cell line representation

Each cell line was originally represented using its untreated gene expression profiles (log-transformed from the Transcripts Per Million (TPM) data) downloaded from DepMap. We further utilized the following embeddings methods to embed each cell line from its original gene expression profiles:

###### Autoencoder embeddings

Given the limited number of cell lines (1,200 in total) with available transcriptomic profiles, training a large-scale foundation model solely on these data would be prone to overfitting and unlikely to yield substantial benefits. Therefore, we considered simpler model architectures for training, such as an autoencoder. The autoencoder was used to generate 128-dimensional cell line embeddings from the original gene expression profiles 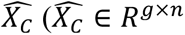, *g* is the number of genes, *n* is the number of total cell lines). Cell lines in the validation sets (for more details on validation sets, see the “**Benchmarking procedures**” section below) were excluded from the autoencoder training, and the rest of the 1,144 cell lines were used for autoencoder training. MSE loss is applied and the objective function is defined as:

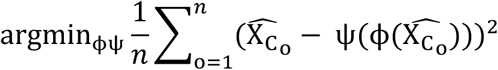

where **φ** and ψ represents the encoder and the decoder, respectively.

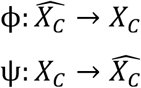

*X_C_* (*X_C_* ∈ *R*^128×*n*^) is extracted as cell line embeddings. The autoencoder was trained for 200 epochs. For cell lines in the test sets, the trained autoencoder was used to directly generate embeddings.

###### scGPT, GeneFormer, and scVI embeddings

Because few pretrained models were available for embedding bulk transcriptomic profiles form cell lines, we considered several SOTA models, including scGPT^68^, GeneFormer^69^, and scVI^70^, originally developed for single-cell transcriptome embeddings. Since we only had around 1,200 cell lines, and the data size was too small to fine-tune these large pretrained models, we opted to use these pretrained SOTA models for zero-shot prediction of the cell line embeddings for benchmarking purposes. To obtain zero-shot embeddings with Geneformer, we employed the 12-layer Geneformer model pretrained on Genecorpus-30M, a large-scale corpus of human single-cell transcriptomes, to generate 512-dimensional cell embeddings. The batch size was set to 10. To generate zero-shot embeddings with scGPT, we loaded the version pretrained on 33 million healthy human cells from the CELLxGENE collection. The model comprises 12 transformer blocks, each with eight attention heads and a hidden size of 512, yielding 512-dimensional cell embeddings. The batch size was set to 64. To obtain scVI embeddings, we first assigned all cells to the same batch label, applied log-normalization, and selected the top 2,000 highly variable genes using Scanpy. We then trained the scVI model with its default architecture, comprising two fully connected layers in both the encoder and the decoder, with a 16- (or 128-) dimensional latent space. Training was performed with a mini-batch size of 128 samples. As reported in Results sections, none of them outperformed the autoencoder approach (Figure S3-S4), which could be due to the discrepancy between their pretrained domain of single-cell transcriptome and the target domain of cell line transcriptome, thus we did not pursue other foundation models further.

##### (3) Protein / Transcription factor representation

Protein and transcriptional factor embeddings were generated using the ProtT5-XL-UniRef50 encoder model, a protein language model based on the T5-3B architecture and pretrained in a self-supervised fashion on a large corpus of protein sequences from UniRef50^71^. The model learns contextualized amino acid representations from large-scale protein sequence data and produces fixed-length embeddings suitable for downstream predictive tasks. Input protein sequences were preprocessed by replacing ambiguous or uncommon residues (U, Z, O, B) with X and inserting spaces between amino acids to match the tokenizer input format. The sequences were truncated to a maximum length of 1,200 residues to reduce computational cost while retaining most structural information. For each sequence, the model produced a per-residue embedding up to the maximum sequence length and a mean pooling was computed to obtain a single fixed-length per-protein embedding vector. These embeddings were used as input features for downstream model training and prediction. All inference was performed on GPUs using half-precision (FP16) computation to improve memory efficiency.

##### (4) Gene representation

We generated gene embeddings using Gene Ontology (GO)-based graph representations. First, GO annotations for all genes were obtained using the ontologySimilarity package^72^, resulting in annotations for 19,614 genes. A bipartite gene-GO graph was then constructed in which genes and GO terms were represented as nodes, and edges connected genes to their associated GO terms. To derive low-dimensional representations of genes, we applied the node2vec algorithm implemented in the PecanPy package^73^. Node2vec learns node embeddings by performing biased random walks on the graph and optimizing node representations to preserve local and global graph structure. Using this approach, 128-dimensional embeddings were generated for all nodes in the gene-GO graph. The objective function of node2vec is defined as follows.

Let *G* denote the set of all genes (|*G*| = 19614) and *T* the set of GO terms. Construct a bipartite graph ℬ = (V, ℰ) with

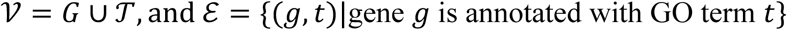

We applied the node2vec algorithm to ℬ. For a node *v*, the transition probability to a neighbor *x* in a random walk is defined as

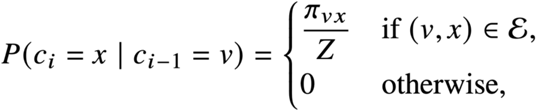

where *π_vx_* = *α_pq_*(*t*, *x*) ⋅ *w_vx_* is the unnormalized transition probability, *w_vx_* is the edge weight (here *w_vx_* = 1 for all edges), and *Z* is the normalizing constant. The bias *α_pq_*(*t*, *x*) depends on the relationship between the previous node *t*, the current node *v*, and the candidate neighbor *x*, and is governed by parameters *p* (return parameter) and *q* (in-out parameter). Node2vec then learns a mapping *f*: V → *R*^128^ that maximizes the log-probability of observing network neighborhoods using a skip-gram architecture. The resulting embeddings {*f*(*g*)}_g∈*G*_ ⊂ *R*^128^ serve as the final gene embeddings.

#### Multi-task training

InsilicoCell stage 1 training is multi-task training, where combined training set data from seven different tasks were used (see the “**Cellular functional profiles**” section). Each input sample for each task is a sequence of multi-modal token embeddings (Figure S1B). During training, every training batch contained a subset of data from all seven tasks, where the same fraction (1/5000 in our experiments) of samples were taken from each of the seven tasks. The released InsilicoCell model contains over 46 million hyperparameters and was trained for 100 epochs on three NVIDIA H100 GPUs for 3 days.

##### Tokenization networks

The tokenization networks are neural network layers which learn a function *f* to transform the original embeddings which have different dimensions into consistent 768-dimensional embeddings *Z*. Each type of original input embeddings has a specific tokenization network, where the transformation can be notated as:

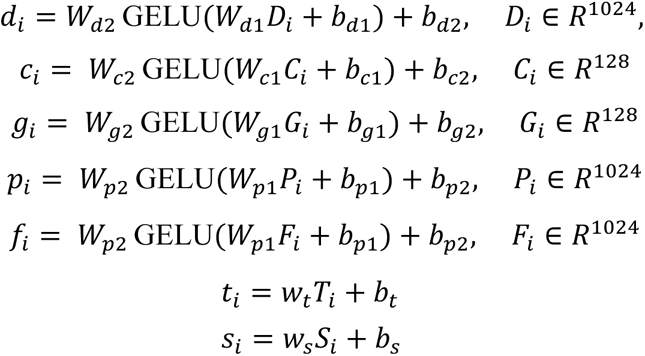

where *D_i_*, *C_i_*, *G_i_*, *P_i_*, *F_i_* are the representations for the inputs of the *i*th labeled data point, as aforementioned in the sections “**Cellular functional profiles**” and “**Input embeddings**”. *W_d_*_1_ ∈ *R*^512×1024^, *W_d2_* ∈ *R*^768×512^, *W_c1_* ∈ *R*^384×128^, *W_c2_* ∈ *R*^768×384^, *W_g1_* ∈ *R*^384×128^, *W_g2_* ∈ *R*^768×384^, *W_p1_* ∈ *R*^512×1024^, *W_p2_* ∈ *R*^768×512^, *w_t_*, *w_s_* ∈ *R*^768^. Note that transcription factors are also a type of proteins, therefore, *P_i_* and *F_i_* use the same set of tokenization networks of *W_p_*_1_, *W_p_*_2_, *b_p_*_1_, and *b_p_*_2_. *T_i_* and *S_i_* are scalar values of the original time and dosage (normalized to [0, 1]), and their tokenization networks make a linear projection. We took reference from Zhou HY et al. for this tokenization approach of scalar values such as time and dosage^74^. Ultimately, the representations of entities in each modality (drug, cell, gene, protein/transcription factor, time, dosage), originally in different dimensions, were converted to token embeddings *d_i_*, *c_i_*, *g_i_*, *p_i_*, *f_i_*, *t_i_*, *s_i_* of the same dimension of 768.

##### Transformer blocks

InsilicoCell contains six transformer encoder blocks with 16 attention heads for self-attention computation. Each input sample is a sequence of token embeddings *Z* (Z ∈ *R^L^*^×768^, L is the sequence length. For multi-task training, L = 6) with the learnable embedding of a class token *CLS* appended at the beginning. Each input sample could contain different types of tokens *Z* depending on the cellular functional profile the sample corresponds to (Figure S1B, and also described in the previous “**Cellular functional profiles**” section), and *PAD* tokens were appended where we needed to make all samples from various tasks have consistent sequence length during multi-task training. Biologically, the cellular functional profiles are not affected by the order of tokens in each input sample (e.g., for predicting drug sensitivity of erlotinib in MCF7, inputting [erlotinib, MCF7] and inputting [MCF7, erlotinib] are biologically the same), therefore, we excluded positional encodings from model design, and InsilicoCell is permutation invariant regarding input samples. For encoder block *b* (*b* ∈ {1,2,…,6}), multi-head self-attention is computed:

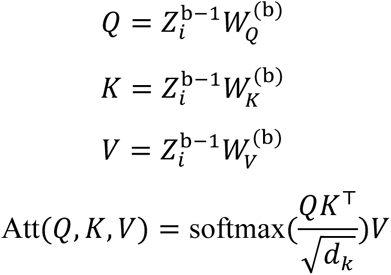

Where *Z_i_* is the sequence of token embeddings (with CLS token appended) for the *i*th sample. W_Q_, W_k_, W_V_ are learnable weight matrices, and *d_k_* is the input embedding dimension per attention head. The feed-forward network has one hidden layer of size 3072 and GELU activation.

##### Prediction heads

The last hidden state of the *CLS* token 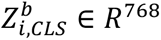 represents the comprehensive information summarized from all the other tokens, which is extracted and passed through a pooler:

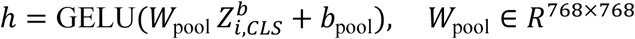

Seven prediction heads are appended to the *CLS* token output, where each head predicts the final result value for one task. Among the seven tasks (*k* ∈ {1,2,…,7}), drug-induced gene expression change, ligand-target binding affinity, drug sensitivity, gene effect score, and copy number variation are regression tasks with continuous values as labels, thus, we used regression heads as prediction heads for input samples coming from these five tasks:

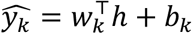

where *w_k_*, ℎ ∈ *R*^768^. While the TF-target gene association and gene mutation are two classification tasks with binary values as labels, thus, we used classification heads as prediction heads for input samples coming from these two tasks:

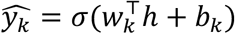

where *σ* represents the logistic sigmoid function.

##### Dynamic loss weighting

During the multi-task pretraining, a dynamic loss weighting method was implemented to calculate total loss according to task difficulty. For each input sample *i*, the original loss *l* is computed between the output from the model’s prediction head 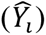 and the ground truth label (*Y_i_*). For the five regression tasks, their original losses *l* are computed using mean squared error (MSE) loss. While for the two classification tasks, their original losses are computed using binary cross entropy (BCE) loss:

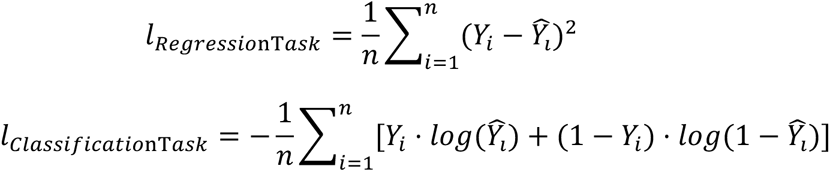

The total loss is computed by weighting each task loss and summing them up. We designed the following dynamic loss weighting method to adjust task weights according to task difficulty reflected by the relative rate of loss changing. This method combines dynamic loss averaging (DWA)^75^ together with task loss normalization. Our dynamic loss weighting method can be notated as:

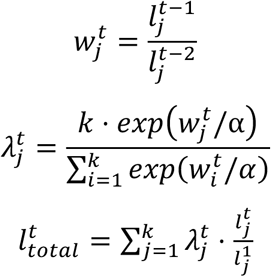

where 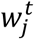 represents the relative loss decreasing rate corresponding to the *jt*ℎ tasks (*j* ∈ {1,2,…,7}) at training epoch *t* (*t* > 2), and 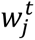 is calculated as the ratio of the task loss *l*_j_ at epoch *t*-1 to that at previous epoch *t*-2. As a result, tasks with slower loss improvement obtain larger values for *w*. *k* represents the total number of tasks, which is 7 here. *α* is a parameter controlling the softness of task weighting, and we empirically set 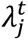 represents the weight for the *jt*ℎ task at training epoch *t*. Consequently, larger weights are assigned for tasks with slower loss improvement. For total loss calculation, in order to prevent the negative effects where some tasks have extremely different loss magnitudes and only the tasks with large loss magnitudes dominate model training, we normalized each task loss using its initial loss in the first training epoch, 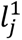 (averaged across all batches in the first epoch), which takes all task losses to similar magnitudes. The weighted normalized task losses are summed together as the total loss 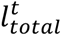. Our validation results showed that the dynamic loss weighting method helped improve InsilicoCell’s prediction performance compared to merely summing up all tasks’ losses as the total loss during multi-task training (Figure S3).

#### Task-specific training

InsilicoCell stage 2 training is task-specific training. Trained model weights from the stage 1 multi-task training were used as the initial weights for stage 2 training. The model was trained seven times, resulting in seven separate models. Each time we only used the training set data of one single task to train one model. The number of epochs for training each task was determined by early stopping based on an internal validation set consisting of 10% randomly holdout samples from the training set for the corresponding task. The task-specific training computes *l*_*Regressio*nT*ask*_ and *l_Classificatio_*_nT*ask*_ depending on the task type, as described in the previous sections. After stage 2 training, each of the seven models can be used for the corresponding task prediction.

#### Active learning (AL)-based few-shot adaptation

For supervised domain adaptation to patient-level prediction (e.g., predict drug sensitivity), InsilicoCell supports both the conventional supervised fine-tuning approach based on the labeled data provided by users and an active learning-based framework for selecting a pre-defined number of optimal samples for subsequent manual label acquisition (e.g., through experiments) and few-shot supervised fine-tuning. We evaluated various active learning algorithms in our previous study^53^. Based on the computation efficiency and performance, as well as the nature of the drug sensitivity task as a regression problem, embedding-based (e.g., based on K-means sampling) active learning is an optimal approach. We modified and adapted the active learning-based sample selection algorithm to the InsilicoCell model as below:

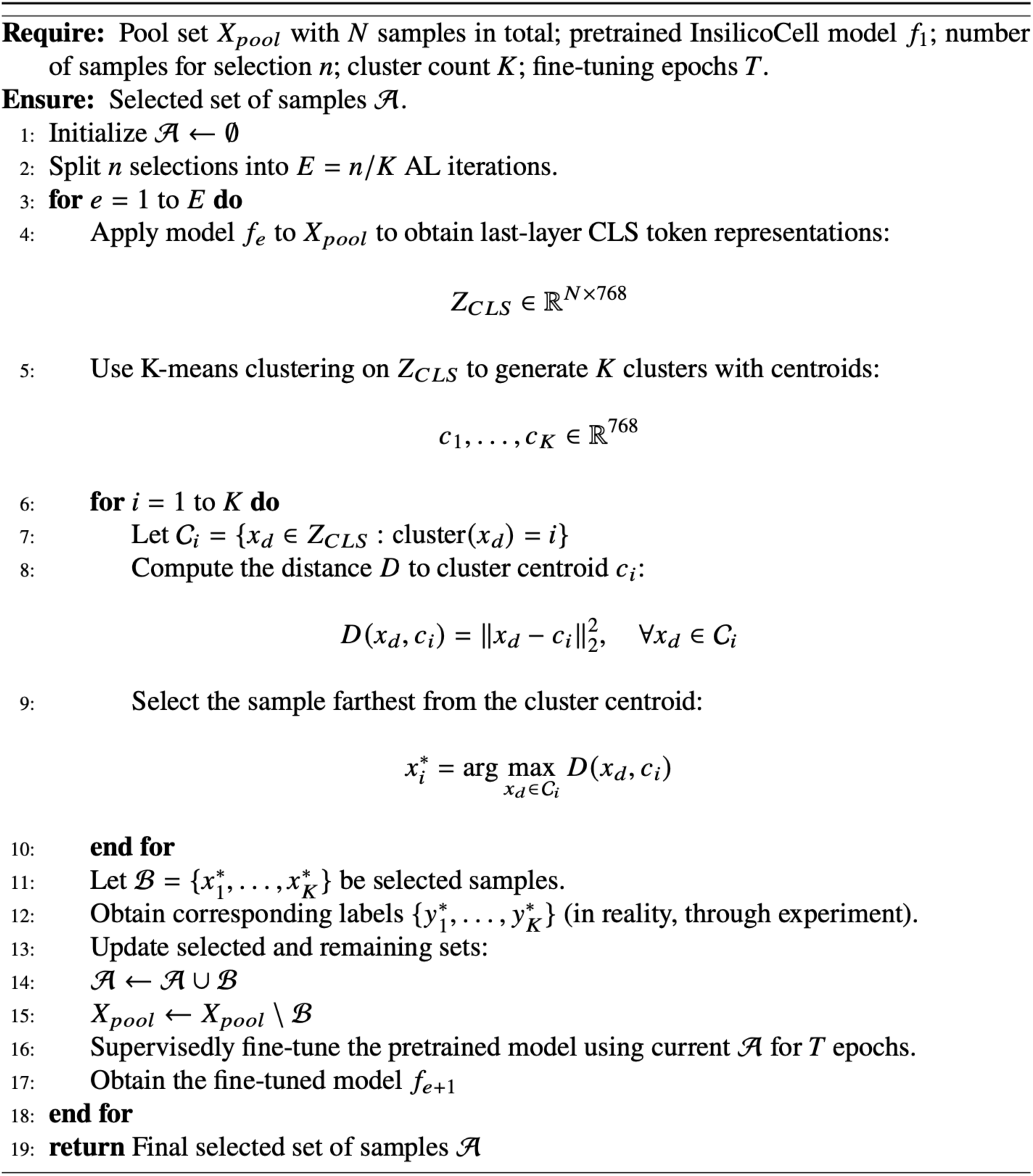

In our setting for predicting drug sensitivity in AML patients, for each patient, 30% of the drug-patient pairs, together with their corresponding drug sensitivity labels, were randomly held out as the test set, and the rest of the data served as an initial pool for selecting few-shot data for model fine-tuning. *X_pool_* represents all the drug-patient pairs initially available. We evaluated multiple selection size (*n* ∈ 300,400,…,1000) with *K* = 100 and *T* = 10. The final selected *n* drug-patient pairs *A* and their ground truth drug sensitivity labels were used for supervised fine-tuning the model *f*_1_ for 10 epochs. We repeated both the active learning-based sample selection as well as random sample selection (baseline for performance comparison) for three runs using three different random seeds, and reported the average performance across all random seeds.

#### Benchmarking procedures

Two different scenarios were considered for prediction performance evaluation: sample-level holdout validation and entity-level holdout validation. Performance of InsilicoCell was compared to various baseline models among all seven tasks. The two tasks of drug-protein binding and TF-gene association were evaluated based on protein/TF-wise performance, and the other five tasks were evaluated based on cell-wise performance. To ensure robust performance estimates, each protein/TF/cell line should contain at least 30 labeled samples, and entities with too few labeled samples were excluded for evaluation. Regarding the evaluation metrics, the five regression tasks, including drug-induced gene expression change, drug-protein binding affinity, drug sensitivity, gene effect score, and CNV, were evaluated using Pearson correlation (*r*) and root mean square error (RMSE) between the prediction and ground truth by default, unless specified otherwise. The other two classification tasks, including TF-gene association and gene mutation, were evaluated using area under the receiver operating characteristic curve (AUROC) and macro F1 score between the prediction and ground truth by default, unless specified otherwise.

##### Entity-level holdout validation

This scenario was used for evaluating model prediction performance on input samples containing previously unseen entities, such as unseen cell lines and unseen compounds, which did not appear in the training set and were viewed by the model as new cell lines and new compounds in the test set to predict on. To construct the entity-level holdout test sets, we first pooled data from all seven tasks and compiled a list of all unique drugs, genes, proteins and cell lines. We then randomly selected 5% of entities from each category. All input sample containing the selected entity were excluded from model training and reserved exclusively for testing. These set-aside samples solely served as test sets for model prediction and performance evaluation. Depending on the specific type of set-aside entities, the entity-level holdout can be specified throughout this paper as drug-level holdout, gene-level holdout, protein-level holdout or cell-level holdout.

##### Sample-level holdout validation

After excluding input samples for entity-level holdout validation as aforementioned, we also created another scenario to evaluate model performance based on sample-level holdout validation using the rest of the data. This was achieved by randomly selecting 10% of all input samples from the left data, serving as a test set that did not participate in model training. In this scenario, all the input samples in the test set were never seen during training, though the entities in the test set could have appeared in the training set. Accordingly, model performance is generally better under sample-level validation than under the more stringent entity-level validation setting.

After setting aside input samples for all the aforementioned test sets based on sample-level holdout validation and entity-level holdout validation, the rest of the data served as the training set for InsilicoCell.

#### Baseline models

The prediction performance of InsilicoCell on various types of cellular functional profiles were compared to a series of baseline models, including baseline models such as MLP and random forest, as well as other SOTA task-specialized models such as DeepConv-DTI, EnzPred-CPI, ConPLex, DTIAM^12–15^ for drug-protein binding affinity prediction; GPS^8^ and DeepCE^7^ for drug-induced gene expression change prediction; the pretrained model TransCell^4^ for drug sensitivity and gene effect score prediction. For each performance comparison, the same test set was used between InsilicoCell and the baseline model.

##### (1) Multilayer perceptron (MLP)

The MLP served as the first baseline model for performance comparison. The training of MLP followed the same 2-stage training as InsilicoCell. The data used for training and testing were also the same as InsilicoCell. The MLP consists of fully connected (FC) layers with GELU as activation function, and seven task heads. Unlike transformers which allow for varying sequence length as input, MLPs require every input sample to contain the same number of total features. Thus, the input layer contained 2,306 neurons (the maximum number of total input features is 1024 (drug) + 128 (cell line) + 128 (gene) + 1024 (protein) + 1 (time) + 1 (dose) = 2306, when the sample comes from the drug-induced gene expression task). Input samples with fewer than 2306 features were zero-padded to the full input dimension. We designed the number of neurons in the MLP’s FC hidden layers as 8177, 3072, and 768, to make the MLP contain approximately the same number of parameters as InsilicoCell for fair comparison. These FC layers were shared by input samples across all seven tasks. Seven task heads (the same as previously described in the “**Prediction heads**” section) were appended to the last FC hidden layer to generate task-specific outputs.

##### (2) Random forest

The random forest model served as the second baseline models for performance comparison. We implemented XGBoost-based random forest models using the XGBRFRegressor and XGBRFClassifier functions from the XGBoost package. All hyperparameters were set to their default setting. As random forest models do not support the multi-task learning framework, we trained seven separate models using data from each task only, which were the same training data as the corresponding single-task InsilicoCell model.

##### (3) DeepConv-DTI

DeepConv-DTI was included as a baseline model specialized for drug-protein binding prediction. The model represents compounds using Morgan fingerprints and encodes protein sequences with convolutional neural networks to extract local residue patterns. As one of the early deep learning frameworks for DTI prediction, DeepDTI provides a widely adopted benchmark architecture for modeling drug-target interactions. We implemented it following the official DeepConv-DTI repository: https://github.com/GIST-CSBL/DeepConv-DTI, and trained DeepConv-DTI using the same training data from the drug-protein binding affinity task as InsilicoCell.

##### (4) EnzPred-CPI

EnzPred-CPI was included as a baseline model specialized for drug-protein binding prediction. It models enzyme-substrate interactions by combining pretrained protein language model representations with learned molecular embeddings within a compound-protein interaction prediction framework, representing a commonly used paradigm for leveraging foundation protein models in compound-protein interaction prediction. We implemented EnzPred-CPI using the default settings provided in the official EnzPred-CPI repository (https://github.com/samgoldman97/enz-pred), and trained EnzPred-CPI using the same training data from the drug-protein binding affinity task as InsilicoCell.

##### (5) ConPLex

ConPLex, a baseline method specialized for drug-protein binding prediction, models interactions by projecting pretrained protein language model embeddings and molecular fingerprints into a shared latent space and learning interaction similarity through contrastive learning, representing an embedding-based paradigm for compound-protein interaction prediction. We implemented it using the default settings provided in the official ConPLex repository: https://github.com/samsledje/ConPLex, trained the model using the same training data from the drug-protein binding affinity task as InsilicoCell.

##### (6) DTIAM

DTIAM, published in 2025, represents a recent state-of-the-art framework specialized for drug-protein binding prediction. It encodes protein sequences using pretrained protein language models and provides a drug molecular pretraining module based on multi-task self-supervised learning for extracting the features of both individual substructures and the whole compound from massive amounts of the molecular graph. We implemented it using the default settings provided in the official DTIAM repository: https://github.com/CSUBioGroup/DTIAM, trained the model using the same training data from the drug-protein binding affinity task as InsilicoCell.

##### (7) TransCell

TransCell was included as a baseline model for predicting drug sensitivity and gene effect scores solely from the transcriptome of untreated cell lines. Although numerous models have been developed for these tasks, many rely on additional features such as mutations to embed cell lines, making direct comparison difficult in our transcriptome-only setting. We therefore selected TransCell^4^, a recently published framework designed to predict cellular responses from gene expression alone, and generated predictions through its web portal. To avoid data leakage arising from overlap between the evaluation datasets used in this study and the pretraining data of TransCell, we restricted evaluation to cell lines that were unseen by both TransCell and InsilicoCell. For drug sensitivity prediction, we selected 13 unseen cell lines from the CTRP dataset as the test set. For gene effect score prediction, we selected 172 unseen cell lines from the DepMap dataset (release 22Q1) as the test set. Because TransCell outputs normalized drug sensitivity scores, whereas InsilicoCell predicts AUC, the two methods produce predictions on different scales. Therefore, for drug sensitivity prediction, performance was evaluated using Spearman correlation coefficient (*r_s_*) between predicted and observed values.

##### (8) GP

GPS was included as a baseline model specialized for drug-induced gene expression change prediction. GPS is a transcriptomics-based virtual screening platform that prioritizes compounds through reversal of disease-associated gene expression signatures. Its RCL module is designed specifically for predicting drug-induced gene expression change and its GPS4Drugs module computes z-RGES, a quantitative measurement of drug-disease transcriptomic reversal. GPS was originally developed using the fixed cellular contexts comprising four commonly used cell lines: MCF7, PC3, VCAP, and HEPG2, under the treatment condition of 24 hours and 10 uM. In our study, HEPG2 was originally designated as an unseen cell line in the cell-level holdout validation and therefore was excluded from GPS benchmarking because GPS cannot predict on unseen cell lines. For the other three cell lines MCF7, PC3, and VCAP, we trained the RCL module using the same training data and test data as InsilicoCell. As GPS does not support joint training across multiple cell lines, time points, or dosages, we trained four separate models for the three cell lines, and only data under the condition of 24 hours and 10 uM were selected for training. Furthermore, the RCL module trains and predicts categorical gene expression responses (“upregulate”, “downregulate”, “no change”), therefore, we converted continuous z-scores into the three classes following the same procedures as described in the GPS paper^8^, prior to the training. For prediction performance comparison between GPS and InsilicoCell, the continuous predictions of InsilicoCell were also converted into discrete classes using the same procedures as described in the GPS paper^8^, and performance evaluation was assessed using macro F1 score between prediction and ground truth for both GPS and InsilicoCell. z-RGES computation for transcriptomic reversal-based drug screening was performed as described before^8^.

##### (9) DeepCE

DeepCE was included as a baseline model specialized for drug-induced gene expression change prediction. DeepCE employs an attention-based model architecture for predicting drug-induced gene expression change and represents a widely used framework for this task. We trained DeepCE using the same training data as InsilicoCell. Because DeepCE allows for training on multiple dosages and cell lines, but not multiple time points, we selected only data under the time point of 24 hours for training. In addition, because DeepCE lacks an explicit mechanism for extrapolating to unseen cellular contexts, only the test set data from seen cell lines were used for prediction and performance evaluation.

#### Validation of InsilicoCell prediction on external datasets

Although InsilicoCell supports seven prediction tasks, we placed particular emphasis on three drug-related applications: drug sensitivity prediction, drug-protein binding affinity prediction, and drug-induced gene expression prediction, given their central importance in drug discovery. Accordingly, we conducted additional validation using independent external datasets to rigorously evaluate model generalizability and real-world performance across these three tasks.

##### Evaluation of InsilicoCell drug sensitivity prediction on GDSC

Pharmacogenomic and transcriptomic data were obtained from the GDSC^2^ resource through the PharmacoGx R package (GDSC 2020, version 2-8.2)^32,33^. For each cell line, gene expression profiles were summarized by taking the median expression value across all available samples. Drug sensitivity measurements, represented as AAC (area above the dose-response curve), were summarized in the same manner. Only cell lines present in both the gene expression and drug sensitivity datasets were retained for subsequent analyses. TransCell drug sensitivity metadata were restricted to the matched compound names. For InsilicoCell evaluation, the same gene expression profiles were used, and corresponding compound structures were retrieved from PubChem as SMILES strings and provided as model inputs. Because InsilicoCell predicts AUC, whereas TransCell predicts normalized drug sensitivity scores with different scales, direct comparison of prediction values is not appropriate. However, both prediction outputs assign higher values to less sensitive cell lines. To align the ground-truth labels with the prediction scale of both models, AAC values were transformed to (1-AAC) which showed positive correlations with both InsilicoCell and TransCell predictions. Model performance was evaluated using Spearman correlation coefficient (*r_s_*) between predicted and observed values. Because no cell line in this external validation dataset contained at least 30 labeled drug-response observations, the minimum inclusion threshold was relaxed to 10 observations per cell line. Only cell lines meeting this criterion were included in the final evaluation.

##### Evaluation of InsilicoCell drug-protein binding affinity prediction on ChEMBL

To establish a highly reliable benchmark dataset for zero-shot drug-protein binding evaluation, we constructed an external validation dataset from ChEMBL (version 35, https://ftp.ebi.ac.uk/pub/databases/chembl/ChEMBLdb/releases/chembl_35/). We retained those records with IC_50_ measurements reported in nM and with single-protein targets. To maximize annotation reliability, we further restricted the dataset to records with a target confidence score ≥ 9, corresponding to the highest-confidence target assignments in ChEMBL, and to records with src_id = 1, indicating measurements curated from the primary scientific literature. We then retained only drug-protein pairs supported by at least two activity records and with bounded within-pair variation (pIC_50_ range ≤ 1.0), so that each pair had repeated and reasonably consistent experimental support. For each (SMILES - protein sequence) pair, repeated IC_50_ values were aggregated by their median, and the final regression label was defined as log_10_IC_50_ (in nM). Finally, to ensure a strict zero-shot setting, we removed all drug-protein pairs already present in the InsilicoCell drug-protein binding pretraining data, thereby preventing overlap with the external benchmark dataset. All baseline models were trained on the training split of the InsilicoCell drug-protein binding dataset and selected using early stopping on its test split. Proteins with at least 30 labeled observations were selected for performance evaluation.

##### Evaluation of InsilicoCell drug-induced gene expression change prediction on GEOMeta

LINCS, the largest publicly available resource of drug-induced gene expression profiles, was included as the primary dataset for training InsilicoCell. To independently evaluate model performance on this task, we constructed an external benchmark dataset, termed GEOMeta, comprising a diverse collection of drug-treated bulk RNA-seq profiles from the GEO database. Because LINCS was generated using the L1000 probe-based expression profiling platform and the expression change was computed and normalized within the plate, making the data distribution different from those from RNA-seq, we did not incorporate GEOMeta into model pretraining in order to avoid cross-platform domain shift. Instead, GEOMeta was used exclusively for evaluating InsilicoCell under both zero-shot and few-shot adaptation settings.

The metadata harmonization procedures used to curate GEO studies have been described elsewhere (https://github.com/Bin-Chen-Lab/GEOMeta). Briefly, an automated agentic AI tool was developed to annotate GEO samples, followed by manual inspection. After obtaining the curated metadata table, we selected bulk RNA-seq studies available in the ARCHS4 database (human gene v2.5)^76^, which processed raw bulk RNA-seq data in GEO using the same pipeline, and we kept the genes supported by InsilicoCell. Replicates (under the same GEO accession ID, with the same treatment and context) were aggregated by computing the median TPM expression. Drug-treated and untreated (control) transcriptomic profiles were matched based on shared GEO accession ID, disease, organ system, and experimental setting. For each treated-untreated pair of transcriptomic profiles, drug-induced gene expression change was computed as the log2 fold change based on the transcriptomic expression data (TPM), following the same procedure of preprocessing external transcriptomic datasets as described previously^8^. The untreated transcriptomic profiles were retained as cellular context representations for subsequent model input. To maximize data coverage while maintaining sufficient sample sizes, we selected the time points (3h, 4h, 6h, 8h, 12h, 16h, 24h, 48h, 72h, 96h, 168h) and dosages (0.01uM, 0.1uM, 0.3uM, 0.5uM, 1uM, 2uM, 2.5uM, 3uM, 5uM, 10uM, 20uM, 30uM, 50uM, 100uM) for fine-tuning and testing. This processing ultimately yielded 2,624 unique perturbed transcriptomic signatures comprising 953 unique untreated transcriptomic profiles and 813 compounds. Each perturbed transcriptomic signature represents the drug-induced gene expression changes for all 19,216 genes under a certain combination of drug treatment, time duration, dosage, and cellular context.

For external evaluation, we randomly withheld 20% of all compounds and 20% of all perturbed transcriptomic signatures and set them aside as two test sets: one for drug-level holdout validation, and the other for signature-level holdout validation. The remaining data were used for fine-tuning InsilicoCell. Because of the distributional differences between GEO and LINCS, zero-shot prediction on the external dataset yielded suboptimal performance. Thus, InsilicoCell was fine-tuned using the GEOMeta training set prior to evaluation. The number of epochs for fine-tuning was determined by early stopping based on an internal validation set consisting of 10% of perturbed transcriptomic signatures randomly withheld from the training set. The corresponding untreated transcriptomic profiles were used as cellular context representations for model input during fine-tuning and prediction. All the continuous drug-induced gene expression changes were converted to classification labels (“upregulate”, “downregulate”, “no change”), following the same procedures as described in the GPS paper^8^. Macro F1 score and balanced accuracy between prediction and ground truth were used for evaluation.

To estimate prediction reliability at the signature level, we calculated a predicted perturbation magnitude score for each perturbed transcriptomic signature. Specifically, the variance of the predicted gene expression changes across all 19,216 genes was computed for each signature, and the resulting variance values were normalized to the range of 0-1 across all signatures. Higher scores indicate stronger predicted transcriptional perturbations induced by treatment. We observed a strong positive association between the predicted perturbation magnitude score and signature-level prediction accuracy (Figure S6B), suggesting that signatures predicted to exhibit larger transcriptional responses are generally associated with more reliable predictions. We further investigated whether model performance could be improved by focusing on strong-signal signatures. To this end, only signatures exhibiting large transcriptional changes in the ground-truth data (variance > 0.1 or > 0.35 across the 19,216 measured gene expression changes) were retained for model fine-tuning and evaluation. Restricting the analysis to these high-variance perturbation signatures resulted in improved predictive performance (Figure S6D).

#### Virtual drug screening based on the reversal of disease-associated gene expression in COAD and HCC

InsilicoCell was used for predicting drug-induced gene expression change, under either the common cellular contexts used by LINCS and GPS (MCF7, PC3, VCAP, HEPG2), or disease-specific cellular contexts. For COAD, we used four commonly used colon cancer cell lines: HCT116, HT29, SW480, and SW620; for HCC, we used four widely used liver cancer cell lines: HepG2, Hep3B, Huh-7, and SNU398 for presenting disease-specific cellular contexts. We compared the screening performance of InsilicoCell to GPS on two previously published benchmark datasets for COAD and HCC, respectively^35^. The datasets contain compounds labeled active or inactive in the corresponding cancer, according to their experimentally measured drug sensitivity (IC_50_). z-RGES was computed between InsilicoCell (or GPS)-predicted drug-induced gene expression change and disease-associated transcriptomic expression data. The predicted z-RGES was used to classify active and inactive compounds.

#### Multi-objective virtual drug screening in HCC

We used InsilicoCell to virtually screen all 1,772,264 compounds in the Enamine HTS library. For each compound, InsilicoCell predicted drug sensitivity, drug-induced *MYC* expression changes, and c-Myc protein activity in HepG2 cells. We used the following filter for selecting the top predicted candidates for subsequent wet-lab validation: (1) predicted drug-induced *MYC* expression change (LINCS-like z-scores of gene expression change values, ranging from −10 to 10) lower than −1, (2) predicted drug sensitivity (CTRP-like AUC values, ranging from 0 to 30) lower than 10, and (3) predicted c-Myc protein activity inhibition (log_10_IC_50_ at nM) lower than 2. At the time of compound procurement, not all prioritized compounds were available in Enamine’s inventory. These compounds which were out of stock were excluded across all drug discovery projects.

To compare the similarity between selected top predicted candidates and pretrained c-Myc activity inhibitors, which have their actual c-Myc activity inhibition (log_10_IC_50_ at nM) measured in BindingDB to be lower than 2, we computed the Tanimoto similarity of the ECFP representations between each top predicted candidate and each pretrained c-Myc activity inhibitor.

#### Multi-objective drug screening in IPF

We applied InsilicoCell to virtually screen all 4,705,628 compounds in the combined Enamine libraries. The single-cell transcriptomic data of myofibroblasts in IPF were downloaded from GSE135893^42^, and converted to pseudobulk data (TPM format) before used as cellular-context inputs for drug screening. For each compound, drug-induced *COL1A1* / *SMA* / *FN1* gene expression change in myofibroblasts, as well as cytotoxicity in fibroblasts represented by drug sensitivity, were predicted by InsilicoCell. We used the following filter for selecting the top predicted candidates for subsequent wet-lab validation: predicted drug-induced *COL1A1* / *SMA* / *FN1* gene expression change in myofibroblasts to be ranked top 1% among all screened compounds (i.e., downregulate all three genes together the most). Among them, we selected two compounds, Z3489603503 and Z2327235199, with the highest predicted cytotoxicity in fibroblasts, as well as three compounds, Z8413727732, Z8874976349, and Z14765088, with predicted reduced cytotoxicity in fibroblasts for validation.

The dose-response curve for the FN1 inhibition effect of Z8413727732 was generated using the “drc” package in R, using the “drm” function, with the parameter selection of “fct = LL.2(names = c("Slope", "IC50"))”.

#### Multi-objective drug screening in MSCs

We applied InsilicoCell to virtually screen all 4,705,628 compounds in the combined Enamine libraries. The transcriptomic data of MSC lines were downloaded from GSE159410^77^ and served as the input for cellular contexts. Drug-induced *OCT4 / SOX2 / NANOG* gene expression change and cytotoxicity (AUC) in MSCs were predicted by InsilicoCell for each compound. We used the following filter for selecting the top predicted candidates for subsequent wet-lab validation: predicted drug-induced *OCT4/ SOX2 / NANOG* gene expression change in MSCs to be >1 among all screened compounds (i.e., upregulate all three genes together the most). Among them, we selected five compounds, Z1903071952, Z2964511831, Z9157927006, Z9348865178 and Z4088391964, with predicted lowest cytotoxicity in MSCs for validation.

#### Spatial-level prediction

##### Spatial transcriptomic data preprocessing

Spatial transcriptomic data of ER+ breast cancer and TNBC tissues, as well as the single-cell transcriptomic data were downloaded from the published study^57^. The representative cell type for each spot was annotated using Seurat’s “FindTransferAnchors” and “TransferData” functions^78^, where the single-cell transcriptomics served as the reference. The raw count data were denoised using SAVERX^79^. The denoised counts were converted to log TPM values. Cell line-level AUC values for the two breast cancer subtypes were from the CTRP database (Figure 6B, right).

##### Domain adaptation to spatial transcriptomes

The pre-processed spatial transcriptomic data were used to fine-tune the autoencoder in the cell line representation module (see the **Cell line representation** section aforementioned) in an unsupervised manner for 200 epochs. The generated embeddings for these spatial transcriptomic data were used as the input of the transformer module, and InsilicoCell made zero-shot prediction of tamoxifen drug sensitivity and tamoxifen-induced gene expression change per spot.

#### Drug combination synergy prediction

Labeled drug combination synergy data^80^ were obtained from GDSC^2^ and used for fine-tuning InsilicoCell (Table S1). For this task, each input sample contains embeddings for two drugs and a cell line, and the prediction outputs a synergy class (whether or not the two drugs have synergic effects in the cell line). To adapt the pretrained InsilicoCell model to the new task, we implemented a transfer learning approach, where the pretrained weights for the multi-modal representation module, the tokenization networks, and the transformer encoder blocks were loaded as the initial weights for fine-tuning InsilicoCell on the new task. A new prediction head (classification head) whose weights were randomly initialized was appended to the CLS pooling output for predicting drug synergy (Figure S12A). We hypothesized that the knowledge learned from the previous seven types of cellular functional profiles (Figure 1A) could improve model performance on other new types of cellular functional profiles as well.

We evaluated InsilicoCell performance on sample-level holdout test set and cell line-level holdout test set, respectively. First, we randomly selected 20% of the unique cell lines from the whole data, and all samples containing these cell lines together served as the cell line-level holdout test set excluded from fine-tuning. Then, from the remaining samples, we randomly selected 10% of them as the sample-level holdout test set. The data excluded the two test sets served as data for fine-tuning InsilicoCell. Macro F1 score and AUROC between prediction and ground truth were used as metrics for evaluating prediction performance. The prediction performance was reported for both fine-tuned InsilicoCell which used transfer learning as aforementioned, and from-scratch trained InsilicoCell which only used drug synergy data alone for training, without any pretrained knowledge from the previous seven types of cellular functional profiles.

#### User-friendly natural language interface of InsilicoCell

We developed a MCP interface which enables users to operate InsilicoCell via natural language conversations within frontier models including Claude and Codex as well as agentic AI tools. User documentation and cases are provided on https://github.com/Bin-Chen-Lab/insilicoCell. This interface allows users to perform prediction of cellular functional profiles and large-scale virtual drug screening by naturally chatting with InsilicoCell, without the need to write command line instructions or manually configure computing environments. Users receive natural language answers within the same conversation window. The MCP service was deployed on an Amazon Web Services (AWS) server and connected to the pretrained InsilicoCell. Claude and Codex serve as conversational clients and workflow orchestrators mapping requests from users to the corresponding InsilicoCell’s prediction task, while neither language model performs the actual computation for predicting cellular functional profiles. All biological predictions are generated by the InsilicoCell on the remote server. Prior to inference, MCP validates inputs, biological identifiers, and the availability of cellular context, estimating the workload scale based on task-specific execution rates. For prediction based on unseen entities, such as unseen cellular contexts, the interface allows users to upload the necessary data file of untreated gene expression in the corresponding cellular contexts, or authorize its retrieval from public sources. Completed result files are made available via temporary download links. A list of APIs designed for InsilicoCell can be viewed at https://github.com/Bin-Chen-Lab/insilicoCell/blob/main/InsilicoCell/API.md.

### Experimental Methods

#### Experimental validation for drug discovery made by InsilicoCell in HCC

##### Crystal violet assay for HepG2 cell viability and IC_50_ determination

HepG2 cells were seeded into 96-well plates at a density of 100 μL per well (1 × 105 cells/mL) in DMEM supplemented with 10% fetal bovine serum and incubated at 37 °C in a humidified atmosphere containing 5% CO₂. Each experimental condition was performed in triplicate wells. After allowing the cells to adhere, cells were treated with the indicated compounds (Z4767240355, Z4969347305, Z2235801986, Z5129787490, and Z2681892181) at various concentrations. Specifically, cells were exposed to compound concentrations of 15.625, 31.25, 62.5, 125, 250, 500, and 1000 nM, as well as 3.125, 6.25, 12.5, 25, 50, and 100 μM, with a final volume of 100 μL per well, and incubated for 48 h. Cells were then stained with crystal violet staining solution (0.125 g crystal violet powder dissolved in 50 mL of 20% methanol) at room temperature for 30 min. After staining, the plates were washed with water and air-dried at room temperature. Subsequently, 100 μL of 30% acetic acid solution was added to each well to solubilize the cell-bound crystal violet. Absorbance was measured at 570 nm using a microplate reader. Cell viability was expressed as a percentage relative to the dimethyl sulfoxide control group (0.1% DMSO), which was defined as 100% viability. Dose-response curves and IC_50_ values were generated using GraphPad Prism 10.

##### Protein extraction and western blot analysis for HepG2 cells

HepG2 cells were treated with Z4767240355 (10 μM), Z4969347305 (10 μM), Z5129787490 (10 μM), Z2235801986 (50nM), or Z2681892181 (500nM) for 24 hours. Cultured cell pellets were lysed in Mammalian Protein Extraction Reagent (Thermo Fisher Scientific) supplemented with phosphatase inhibitors. Protein concentrations were quantified using the Bio-Rad Protein Assay Kit (Bio-Rad, Hercules, CA) with bovine serum albumin (BSA) as the calibration standard. Equal amounts of protein were mixed with Tris-Glycine SDS sample buffer (Life Technologies, Carlsbad, CA) and denatured by boiling. Samples were resolved by SDS–polyacrylamide gel electrophoresis (SDS-PAGE) and transferred onto nitrocellulose membranes (Life Technologies) using electroblotting. Membranes were blocked for 1 h in 5% non-fat dry milk, followed by overnight incubation at 4 °C with Anti-Myc tag antibody [9E10] (Abcam; 1:5000) as primary antibodies. After washing, membranes were incubated with horseradish peroxidase (HRP)-conjugated secondary antibodies (Jackson ImmunoResearch, West Grove, PA; 1:5000 dilution, 30 min). Protein signals were visualized with the SuperSignal West Femto Chemiluminescent Substrate (Pierce, New York, NY).

##### Colony formation assay

HepG2 cells were used to evaluate the effects of drug treatment on long-term cell proliferative capacity. Cells in the logarithmic growth phase were harvested and seeded into 6-well plates at a density of 1,000 cells per well. Each experimental condition was performed in triplicate. After cell attachment, cells were treated with either Z4767240355 (25 uM) or Z4969347305 (25 uM), while the control group received an equivalent volume of DMSO. Cells were then cultured under standard conditions for 14 days, with the culture medium and corresponding treatments refreshed as required. At the end of the incubation period, colonies were fixed and stained with crystal violet. Visible colonies were imaged and counted manually. Colony formation efficiency was calculated based on the number of colonies formed relative to the number of cells initially seeded.

##### Dual-Luciferase Reporter Assay for c-Myc Activity

c-Myc–dependent transcriptional activity was assessed in HepG2 cells using a dual-luciferase reporter assay. Cells were seeded into 96-well plates in antibiotic-free medium and allowed to adhere overnight. When ∼70% confluent, cells were transiently transfected with 1 μl of Reporter plasmid from the Myc Reporter Kit (BPS Bioscience, Catalog #60519) using Lipofectamine 3000 reagent (Thermo Fisher Scientific, Catalog #L3000015). Positive controls were co-transfected with the c-Myc expression vector, while negative controls received the Negative Control Reporter alone. 6 hours after transfection, the medium was replaced with complete culture medium containing 10% FBS and 1% penicillin– streptomycin. Twenty-four hours post-transfection, cells were treated with DMSO or the indicated compounds: Z2681892181 (250 nM), Z2235801986 (62.5 nM), Z4969347305 (100 μM), Z4767240355 (100 μM), and Z5129787490 (100 μM), and incubated for an additional 24 hours. The concentrations for effective compounds were chosen based on their IC_50_ derived from dose-response curves in HepG2 cells. Luciferase activities were measured using a two-step (Firefly & Renilla) luciferase assay system (BPS Bioscience, Catalog #60683). Firefly luciferase was normalized to Renilla luciferase, and relative activity was expressed as fold change compared with DMSO-treated controls.

#### Experimental validation for drug discovery made by InsilicoCell in IPF

##### Cell culture and drug treatment

Human lung fibroblasts were maintained under standard culture conditions at 37°C in a humidified incubator with 5% CO₂. To induce fibrotic activation, cells were stimulated with TGF-β and treated with experimental compounds Z3489603503, Z2327235199, Z8413727732, Z8874976349, or Z14765088. Unless otherwise indicated, compounds were administered at 10 μM for 72 h, with DMSO-treated cells serving as vehicle controls. For dose-response studies, fibroblasts were treated with increasing concentrations of Z8413727732 (0.1, 0.3, 1, 3, and 10 μM) in the presence of TGF-β. Nintedanib and nerandomilast were included as positive controls.

##### Cytotoxicity and morphological assessment

To evaluate compound-associated cytotoxicity, fibroblast morphology and cell adherence were monitored by microscopy following drug treatment. Cells treated with Z3489603503 and Z2327235199 exhibited pronounced cytotoxicity, characterized by loss of cell adherence and altered cellular morphology, consistent with the initial hypothesis. In contrast, Z8413727732, Z8874976349, and Z14765088 did not induce overt cytotoxicity and were therefore selected for subsequent antifibrotic analyses.

##### Protein extraction and western blotting

Cells were lysed in RIPA buffer (MilliporeSigma, R0278) supplemented with protease inhibitors (Thermo Scientific, A32955) and phosphatase inhibitor (A32957), and centrifuged at 12,000 × g for 15 min at 4°C. Supernatants were collected, and protein concentrations were determined using the BCA assay (Thermo Scientific, 23235). Equal amounts of protein were separated by 10% SDS-PAGE (Invitrogen, NW04122BOX) and transferred onto PVDF membranes using the iBlot system (25 V, 6 min). Membranes were blocked with 5% BSA in TBST for 1 h and incubated overnight at 4°C with primary antibodies (Diluted in 2.5% BSA in TBST) against: Fibronectin (FN1) (1:1000), -SMA (ACTA2) (1:2000), COL1A1 (1:1000), Phospho-SMAD3 (pSMAD3) (1:1000), GAPDH (loading control) (1:2000), β-actin (loading control) (1:2000). After washing, membranes were incubated with IRDye 680RD and IRDye 800CW fluorescent secondary antibodies (LI-COR Biosciences) for 1 h at room temperature. Fluorescent signals were detected using the LI-COR Odyssey imaging system, and band intensities were quantified using ImageJ. Target protein signals were normalized to GAPDH or β-actin and expressed as fold change relative to the TGF + DMSO control group. Reagent or resource sources and identifiers are: Anti-Fibronectin (FN1) (MilliporeSigma, Cat#F3648; RRID: AB 476976), Anti-*α*-SMA (MilliporeSigma, Cat#A5228; RRID: AB 262054), Anti-COL1A1 (ABclonal, Cat#A1352; RRID: AB 2760381), Anti-GAPDH (Cell Signaling, Cat#3906; RRID: AB2107303), RIPA buffer (MilliporeSigma, R0278), Protease inhibitor (Thermo Scientific, A32955), BCA protein assay kit (Thermo Scientific, 23235), SDS-PAGE gel (10%) (Invitrogen NW04122BOX), TRIzol reagent (Invitrogen, 15596026), RNeasy Mini Kit (Qiagen, 74106).

##### Dose-response and SMAD signaling assay

To evaluate the effect of Z8413727732 on canonical TGF-β signaling, fibroblasts were stimulated with TGF-β and treated with increasing concentrations of Z8413727732 (0.1–10 μM). Total protein was extracted as described above, and western blot analysis was performed to detect phosphorylated SMAD2/3 (pSMAD2/3). pSMAD2/3 levels were normalized to β-actin and quantified relative to TGF + DMSO controls. Dose-dependent suppression of pSMAD signaling by Z8413727732 was used to assess inhibition of the canonical TGF-β/SMAD pathway.

##### IPF precision-cut lung slice (PCLS) assay

To evaluate the antifibrotic activity of Z8413727732 in a physiologically relevant ex vivo model, PCLS were prepared from human IPF lung tissue obtained from Cowell Health Hospital. IPF lung explant samples were obtained from 3 patients undergoing lung transplantation. Pathologic assessment confirmed the findings of advanced UIP in the IPF subjects. All protocols complied with all relevant ethical regulations approved by local Institutional Review Boards (IRB #: CHW 2017-198); Written informed consent was obtained from all patients. Lung slices were cultured under standard conditions in DMEM supplemented with 0.5% FBS and treated with Z8413727732 for 72 hours. Vehicle-treated slices were included as controls. Z8413727732 was evaluated in six independent PCLS experiments to ensure reproducibility. After treatment, lung slices were collected and lysed for total protein extraction. Protein expression was analyzed by western blot using primary antibodies against fibronectin (FN1), α-smooth muscle actin (α-SMA), collagen type I alpha 1 (COL1A1), beta-actin and GAPDH. GAPDH or beta-actin was used as the housekeeping and loading control for normalization across samples.

#### Experimental validation for drug discovery made by InsilicoCell in MSCs

##### WJMSCs isolation

WJMSCs were isolated as described previously^81,82^. Human umbilical cords (UC) with placenta were collected at the Gift of Life Michigan Donor Care Center. Informed consent from the donors was obtained before getting the placenta. The collection procedures and ethical approval were obtained from the institutional ethics committee (IEC). The tissue was collected in a preservation solution on ice and transported to the Pancreatic Cancer Center, Henry Ford Health, Detroit, Michigan. In sterile condition, UC was cut and removed from placenta. UC was washed using Dulbecco’s phosphate buffered saline (DPBS) containing 1X antibiotic solution (Thermo Fisher Scientific, Catalog #15140122). UC was cut vertically to expose the inside jelly portion. Further, the jelly portion was cut into tiny cubical explant pieces. The explants were kept in 35mm tissue culture dishes in the inverted position for proper attachment. Digestion enzyme solution was added gently, and explants were kept for overnight digestion to ooze out cells in a humidified 5% CO2 incubator at 37 °C. Digestion enzyme solution was prepared by mixing serum-free DMEM and 0.25% trypsin-EDTA solution in 1:1 ratio. Next, 0.5mg/ml of collagenase type I (Fisher Scientific, Catalog #NC9482366) was added to the solution, mixed them well, and filtered using 0.2µm syringe filter. Primocin (2ug/ml) (InvivoGen, Catalog #ant-pm-1) was added to the solution to avoid microbial growth. After overnight digestion, the digested tissues were strained using a 100mm cell strainer. The strained cells were centrifuged for 25min at 2200rpm. After centrifugation, the supernatant was discarded carefully without disturbing the pellet. Fresh media was added to the pellet and plated for cell attachment and growth.

WJMSCs were cultured in 50% of KnockOut™ DMEM/F-12 and 50% of L-WRN^83^ conditioned media supplemented with 10% Fetal Bovine Serum (FBS), 1X GlutaMAX and 1X antibiotic solution. Upon reaching 80-90% confluency, cells were trypsinized using 0.25% trypsin-EDTA solution and sub-cultured, passage number 3 to 6 was used for experimental purposes.

##### Cell treatment

Compounds (Z1903071952, Z2964511831, Z9157927006, Z9348865178, and Z4088391964) were procured from Enamine and a 10mM stock solution was prepared using dimethyl sulfoxide (DMSO). WJMSCs (1×10^5^ cells per well) were seeded in 6-well tissue culture plates and incubated for 48 hrs. After that, we treated the WJMSCs with three compounds at a concentration of 10µM for 48 hrs. 0.1% DMSO-treated WJMSCs were used as control.

##### Quantitative Reverse Transcription Polymerase Chain Reaction (qPCR)

To quantify the expression of pluripotency genes *OCT4*, *SOX2*, and *NANOG*, we performed qPCR. After treatment, WJMSCs were collected in TRIzol™ (Thermo Fisher Scientific, Catalog #15596026) for RNA isolation. Total RNA isolation was performed using Zymo-kit. Then, RNA quality and concentration were measured using Nanodrop. The absorbance at 260/280 of ≥1.8 was considered for cDNA synthesis. Next, 500ng of complementary DNA (cDNA) was synthesized using qScript Ultra SuperMix (Quantabio, Catalog #95217-100) kit as per manufacturer’s instructions. qPCR was performed using a Luna® Universal qPCR Master Mix (New England Biolabs, Catalog #M3003X) and the reaction was carried out in a QuantStudio 6 Pro real-time PCR System (Applied Biosystems). The relative gene expression was calculated using 2^-ΔΔCT^ and beta actin, a housekeeping gene, was used for normalization.

Gene primer sequence are shown below:

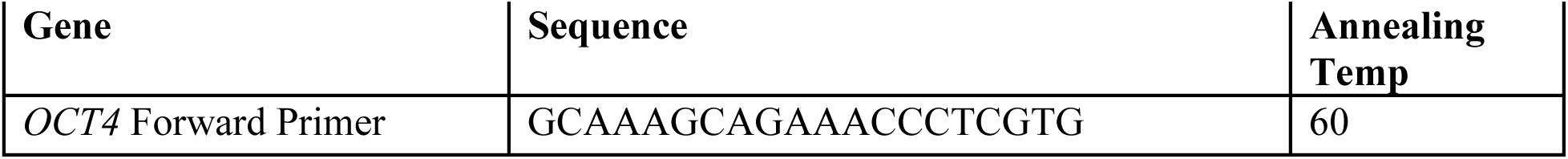
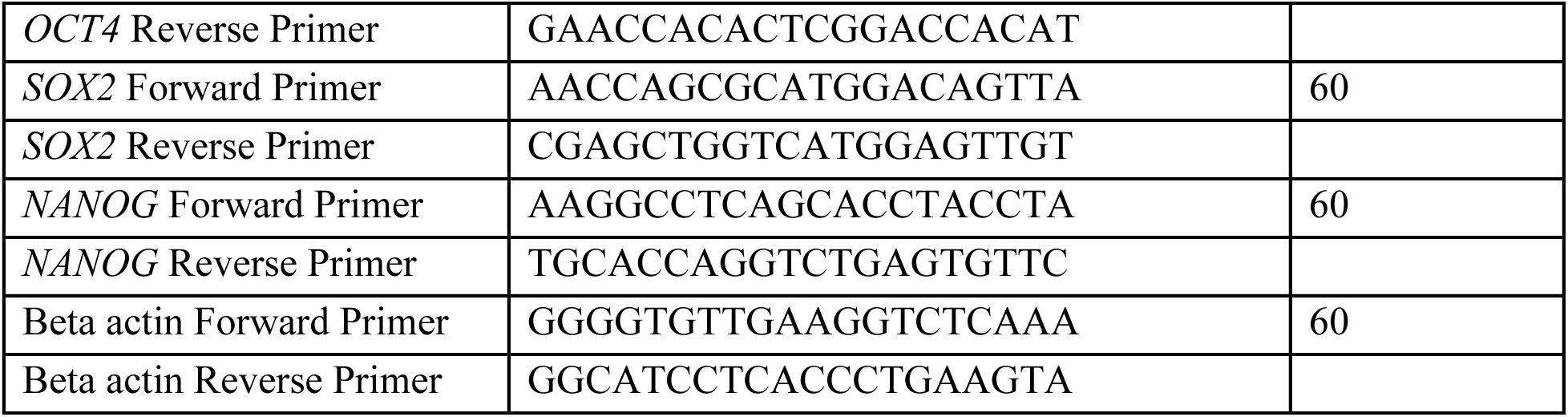

### Statistical analysis

Unless otherwise specified, statistical significance was assessed using two-sided Student’s t-tests. For data that did not satisfy the assumptions of normality, the Wilcoxon rank-sum test was used. For fold change data in western-blot and qPCR experiments, ratio paired t-test was used. Associations between variables were quantified using the Pearson correlation coefficient (*r*) by default. For comparisons involving measurements derived from different populations, experimental platforms, or measurement scales, the Spearman rank correlation coefficient (*r_s_*) was used and explicitly indicated. All statistical tests were two-sided, and P < 0.05 was considered statistically significant.

## Data availability

All datasets are from public sources.

LINCS L5: https://clue.io/releases/data-dashboard.

BindingDB: https://www.bindingdb.org/rwd/bind/chemsearch/marvin/Download.jsp.

CTRP: https://portals.broadinstitute.org/ctrp.v2/.

GDSC^2^: https://gdsc-combinations.depmap.sanger.ac.uk.

ChEA: https://maayanlab.cloud/Harmonizome/dataset/CHEA+Transcription+Factor+Targets.

DepMap: https://depmap.org/portal/.

Patient drug sensitivity data: https://biodev.github.io/BeatAML2/.

Spatial transcriptomic data for breast cancer, scRNA-seq data for IPF, and bulk RNA-seq data for MSCs can be found at National Center for Biotechnology Information Gene Expression Omnibus (GEO) under accession numbers GSE176078, GSE135893, and GSE159410, respectively. The training and test set splitting of InsilicoCell will be available on https://huggingface.co/datasets/binchenlab/InsilicoCell after the acceptance of the paper.

## Code availability

The user documentation, model weights and the code of InsilicoCell are available at https://github.com/Bin-Chen-Lab/insilicoCell.

## Author Contributions

R.C. developed the model, performed benchmarking evaluation and virtual drug screening, and analyzed the results. Y.Q. and X.C. performed bench experiments for the case of HCC. X.L. and L.H. performed bench experiments for the case of IPF. S.M. and L.H. performed bench experiments for the case of MSC. R.C., L.M., and D.L. contributed to baseline comparison. L.M., L.L., J.P., and Y.X. contributed to representation learning. X.Z. provided labeled GEO data. E.E. provided medicinal chemistry support and R.G. provided clinical samples. R.C., B.C., and J.Z. wrote the manuscript with input from all authors. J.Z. supervised model development. B.C. conceived and supervised the study.

## Declaration of interests

B.C. is a co-founder of Bronto Therapeutics. A provisional patent application covering these novel compounds has been filed (B.C., R.C., X.L., and L.H.). Other authors declare no competing interests.

## Acknowledgements

The research is supported by the NIH R01GM134307 (B.C., and J.Z.), R01GM145700 (B.C., J.Z., and E.E.), R61HL177451 (X.L., R.G. and B.C.), the NSF IIS-2212174 (J.Z.), and the MSU SPG grant (B.C., J.Z., E.E., X.L.). We extend our sincere appreciation to the Corewell Health Lung Transplant Program and Dave Chesla, Tara Jager, Cameron Lawson for the management of the Spectrum Health Accelerator of Research Excellence (SHARE) protocol, which facilitates the collection of de-identified human donor lungs.

## Extended data

**Figure S1.**
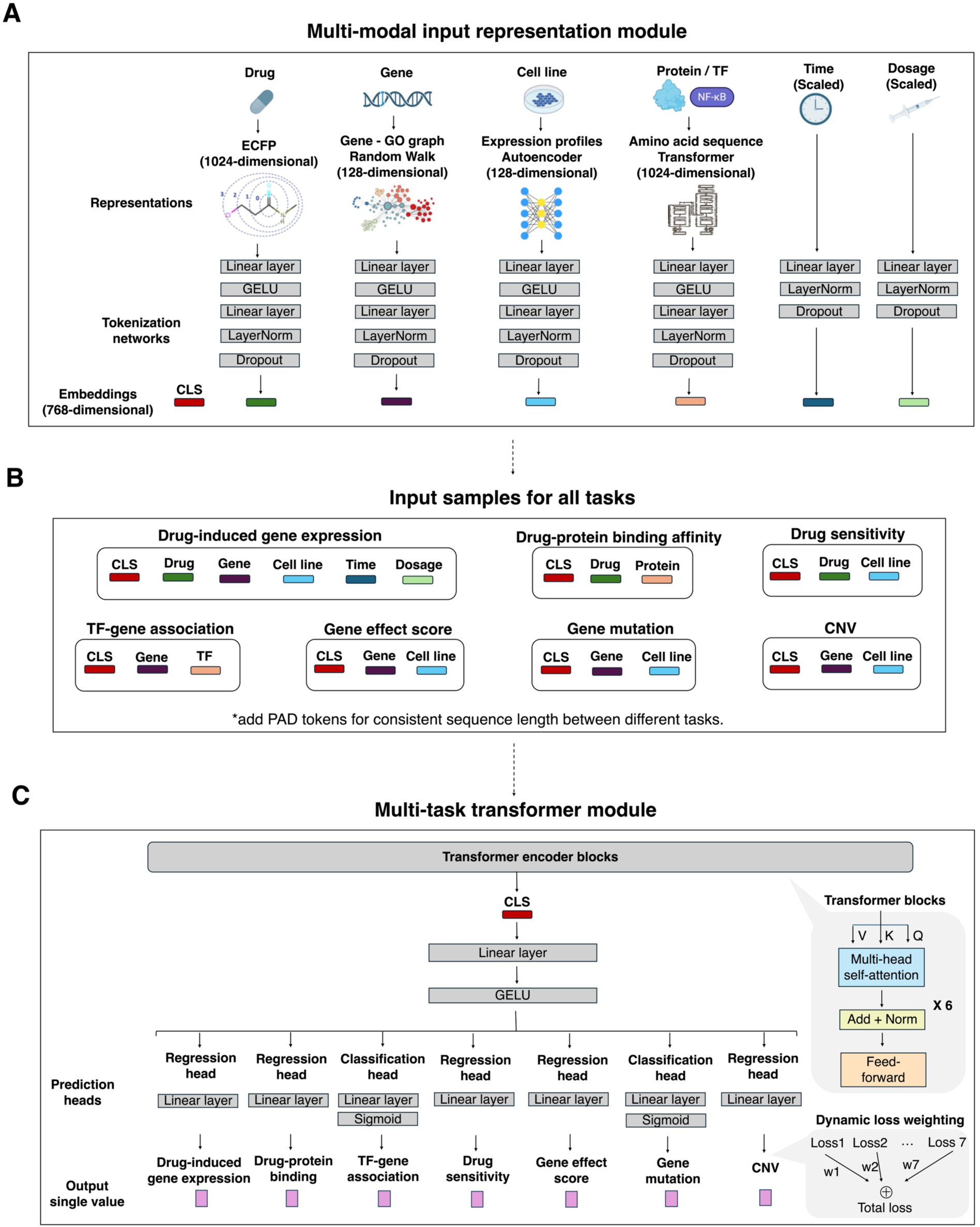
Detailed model architecture of InsilicoCell, Related to Figure 1. **A**, The multi-modal representation module of InsilicoCell. Drugs are represented via ECFP features, genes are represented via a random walk algorithm based on Gene Ontology graph, cells are represented via the latent embeddings from an autoencoder based on the original cellular gene expression profiles, and proteins/TFs are represented via the embeddings from a transformer based on amino acid sequences. Details on generating these representations as well as other alternative representations are listed in Methods. Time and dosage are scaled values. A series of tokenization networks convert all these multi-modal representations into embeddings of consistent dimensions. **B**, Input samples for all seven tasks. Each input sample for each task is a sequence (permutation invariant) of multi-modal token embeddings. For multi-task training, samples with sequence lengths of < 6 are appended with PAD token embeddings. **C**, The multi-task transformer module for label prediction. There are six transformer encoder blocks with multi-head self-attention, residual connection, layer normalization, and feed-forward layers. Hidden states for the CLS token embedding are pooled and appended with seven task-specific prediction heads, with regression heads for five regression tasks with continuous labels, and classification heads for two classification tasks with binary labels. Dynamic loss weighting is used for loss computation in multi-task training stage (Methods).

**Figure S2.**
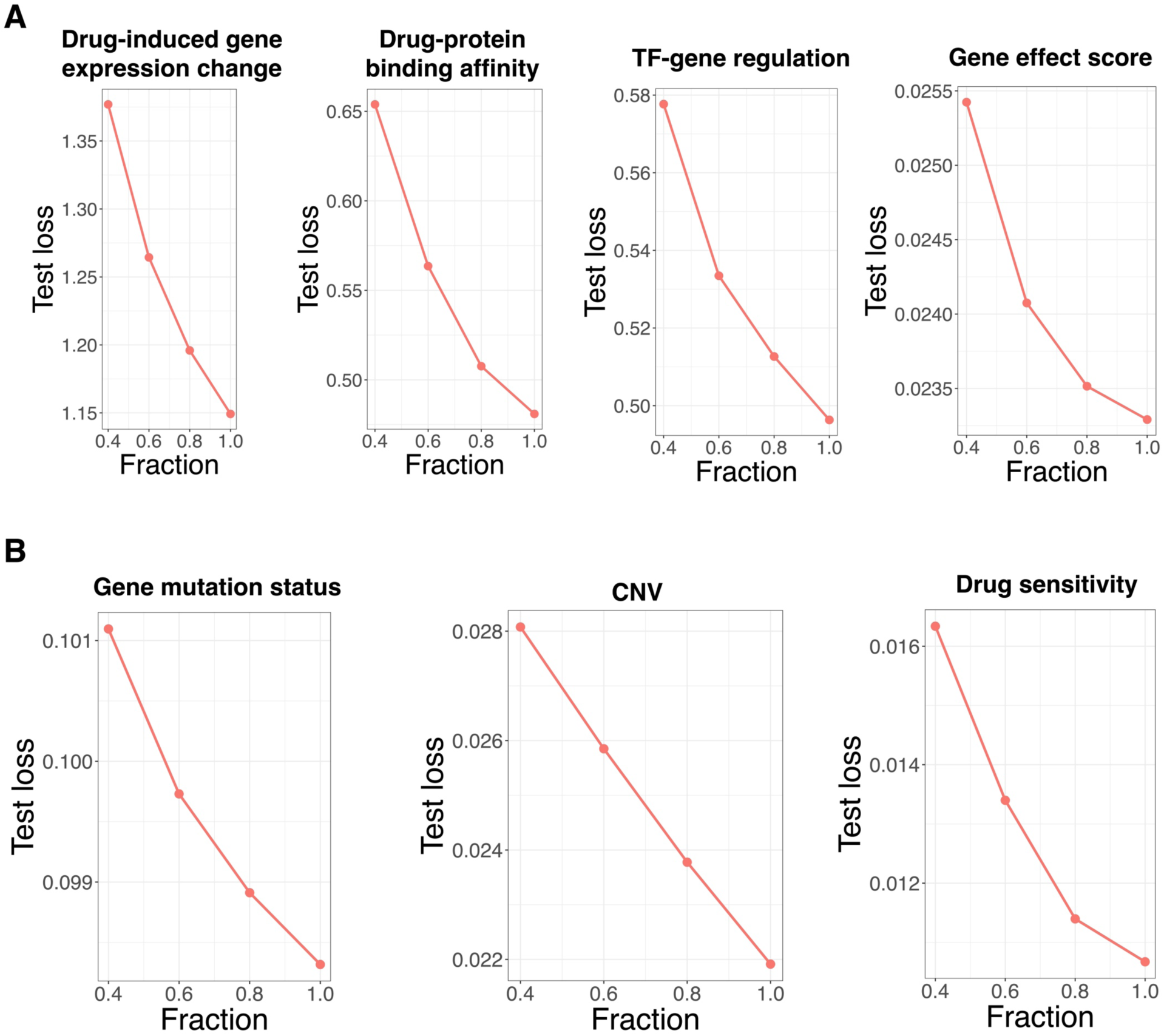
Scalability of InsilicoCell with respect to training input sample size, related to Figure 2. **A-B,** Loss on the test set with respect to different fractions of training input samples (40%, 60%, 80%, 100%). InsilicoCell was trained from scratch on different sample sizes, and the test loss from the multi-task training (stage 1 from Figure 2A) was observed.

**Figure S3.**
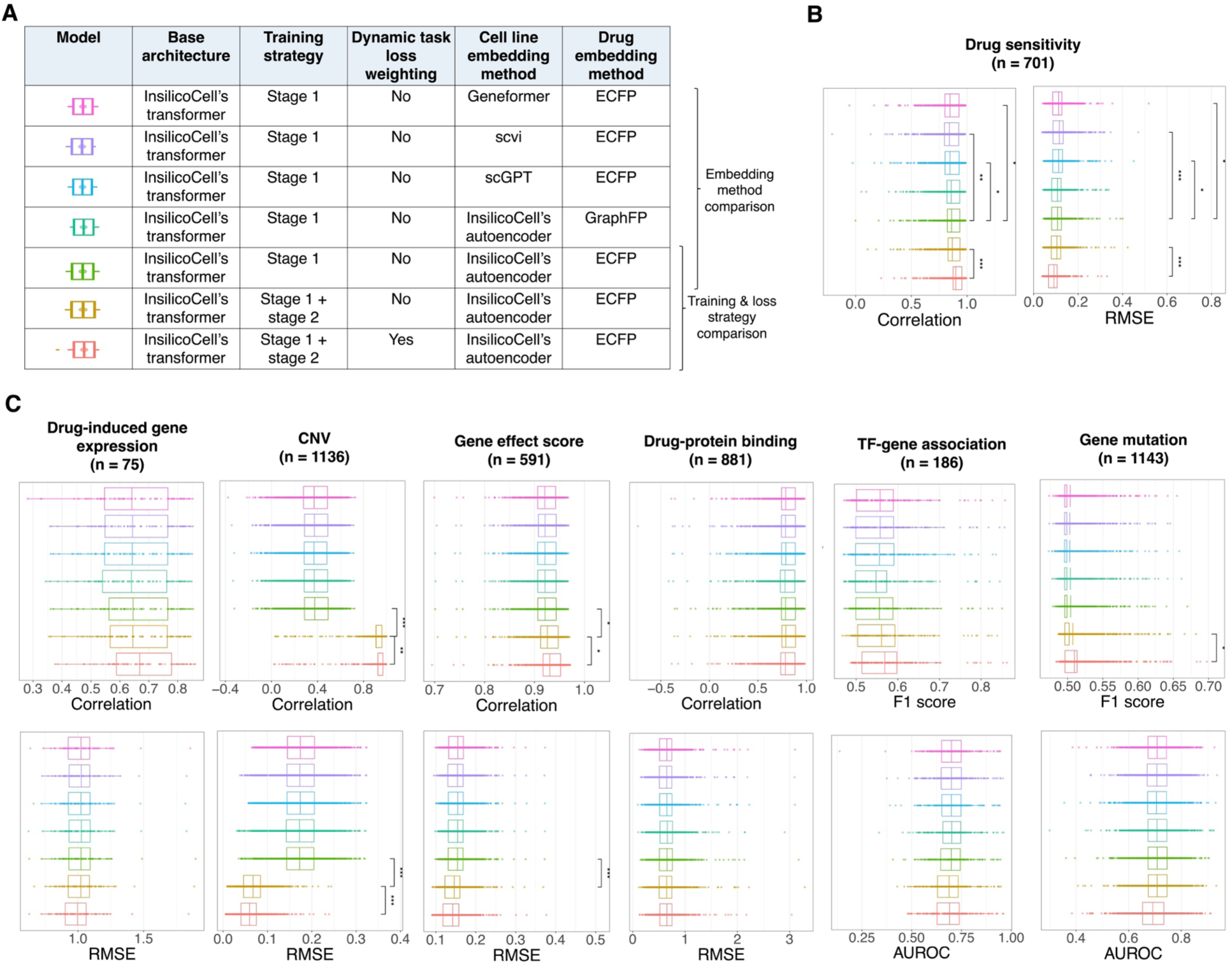
Additional results for the ablation study on the sample-level holdout test set, related to Figure 2. **A,** Details of each baseline model. **B-C.** Boxplots showing prediction performance on the sample-level holdout test set for each cellular functional profile. For drug-protein binding affinity and TF-gene association tasks, the error bars show protein/TF-wise performance. *n* represents the number of proteins. For all the other tasks, the error bars show cell-wise performance. *n* represents the number of cell lines. All five regression tasks use correlation and RMSE between prediction and ground truth as evaluation metrics, and the other two classification tasks use F1 score and AUROC as evaluation metrics. Asterisks mean statistically significant difference via t-test (*: < 0.05, ** < 0.01, *** < 0.001).

**Figure S4.**
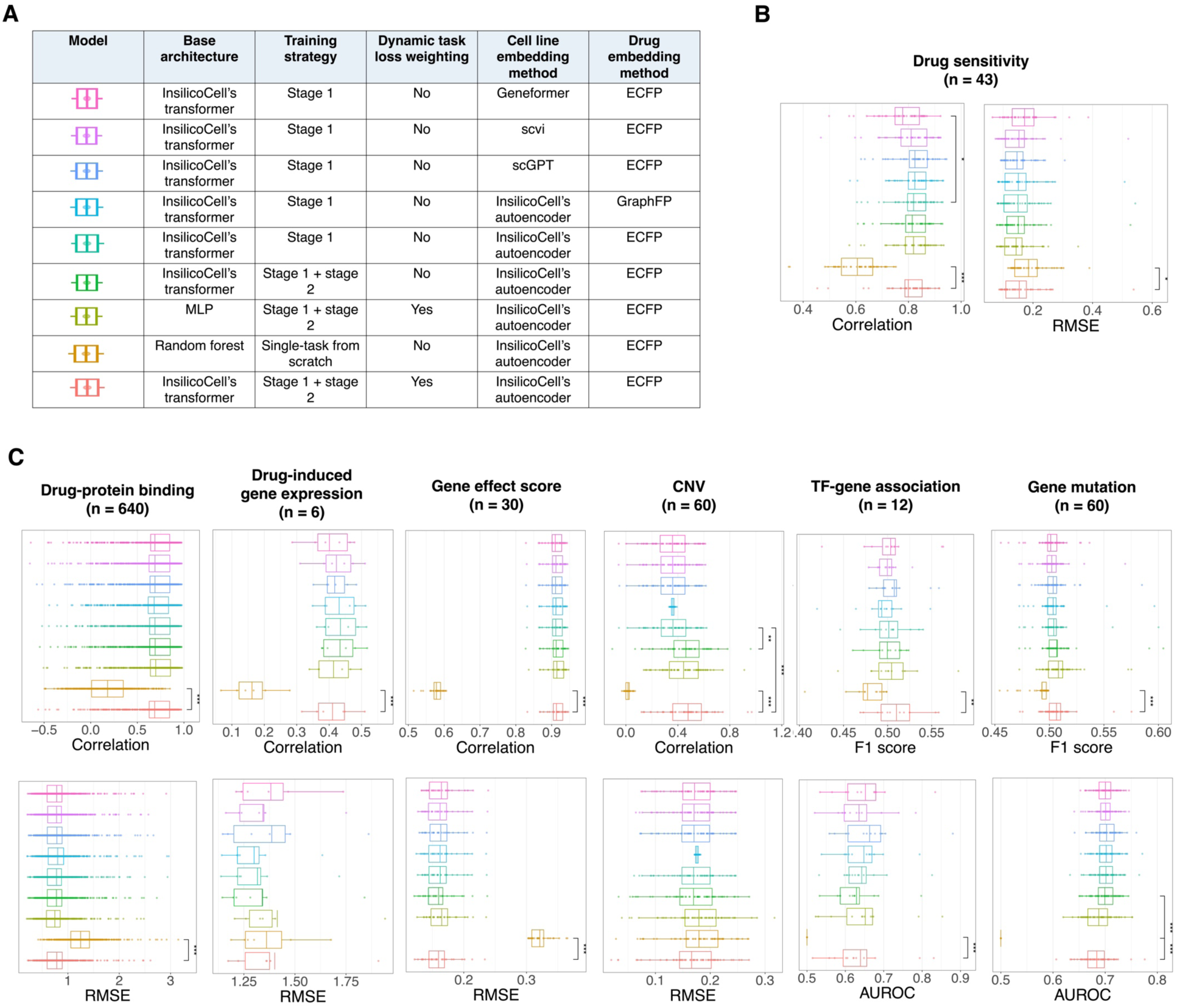
Additional results for the ablation study on the entity-level holdout test set, related to Figure 2. **A,** Details of each baseline model. **B-C.** Boxplots showing prediction performance on the entity-level holdout test set for each cellular functional profile. For drug-protein binding affinity task, drug-level holdout validation was used, where all drugs in the test set were unseen in the pretraining data. The error bars show protein-wise performance. *n* represents the number of proteins. For the TF-gene association task, TF-level holdout validation was used, where all TFs in the test set were unseen in the pretraining data. The error bars show TF-wise performance. *n* represents the number of TFs. For all the other tasks, cell line-level holdout validation was used, where all cell lines in the test set were unseen in the pretraining data. The error bars show cell-wise performance. *n* represents the number of cell lines. All five regression tasks use correlation and RMSE between prediction and ground truth as evaluation metrics, and the other two classification tasks use F1 score and AUROC as evaluation metrics. Asterisks mean statistically significant difference via t-test (*: < 0.05, ** < 0.01, *** < 0.001).

**Figure S5.**
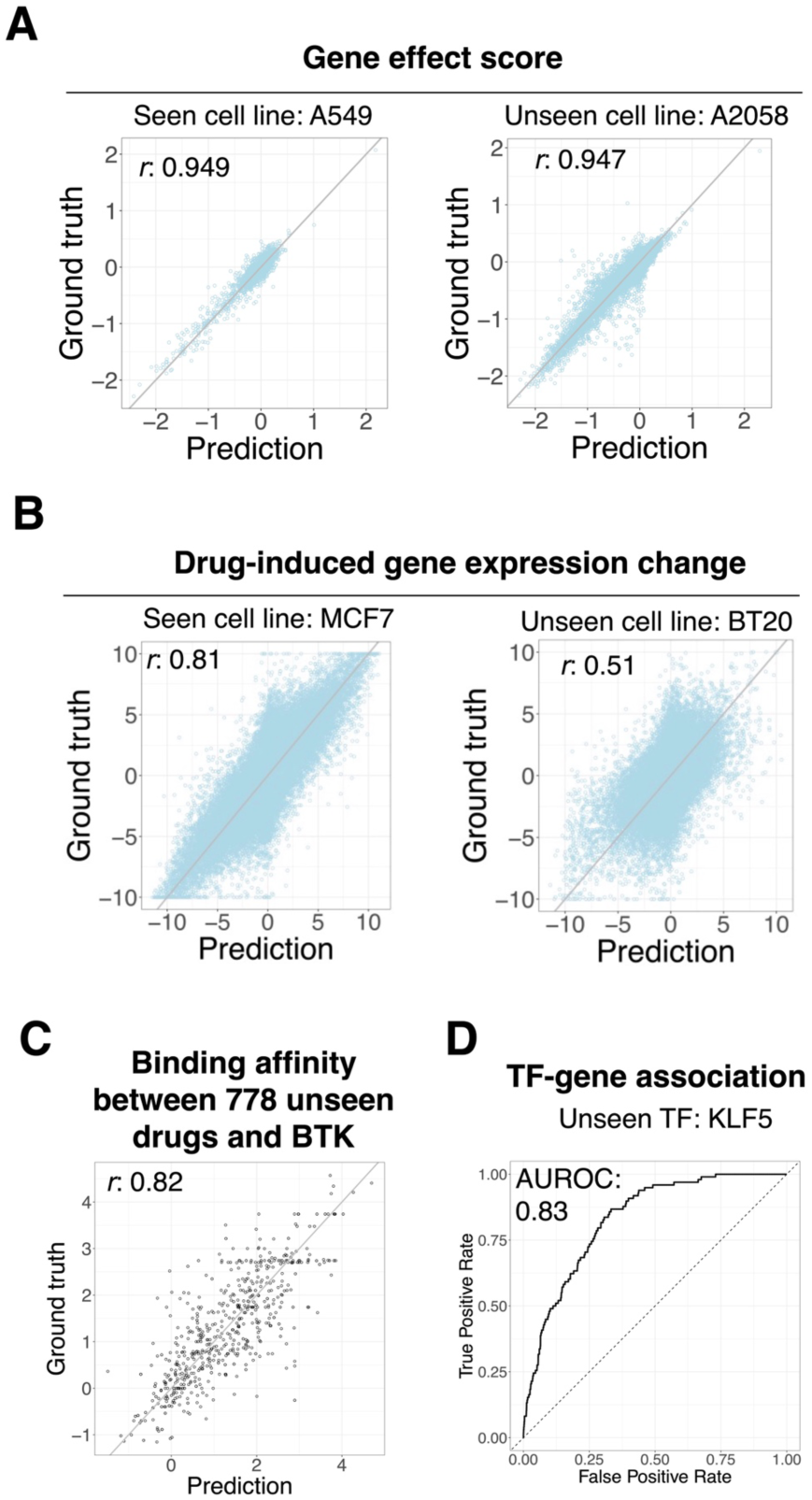
Representative examples of prediction with seen and unseen entities by InsilicoCell, related to Figure 2. **A,** Scatter plots showing the prediction of gene effect scores in a seen cell line A549 (left) and an unseen cell line A2058 (right), where the correlation between prediction and ground truth are 0.949 and 0.947, respectively. **B,** Scatter plots showing the prediction of drug-induced gene expression change in a seen cell line MCF7 (left) and an unseen cell line BT20 (right), where the correlation between prediction and ground truth are 0.81 and 0.51, respectively. **C**, Scatter plot showing the prediction of drug-protein binding affinity for an example protein BTK on the drug-level holdout test set (778 unseen drugs for this protein). The correlation between prediction and ground truth is 0.82. **D**, AUROC curve showing the prediction of TF-gene association between an example unseen TF (KLF5) and all genes.

**Figure S6.**
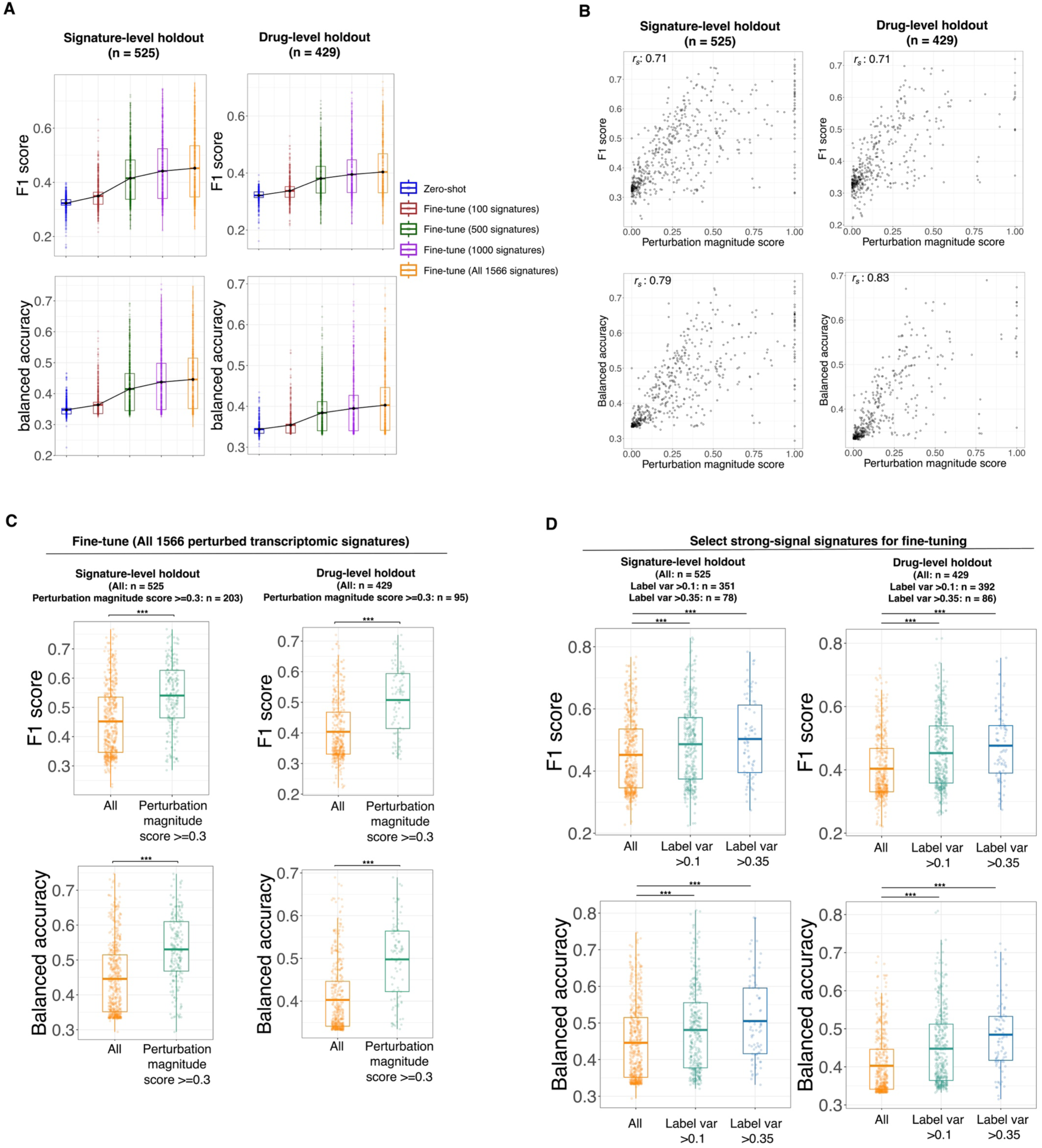
Performance evaluation of InsilicoCell for drug-induced gene expression change prediction with an external dataset of GEOMeta, related to Figure 3. **A,** Performance on the test set based on different numbers of samples randomly selected for fine-tuning as well as zero-shot prediction. For signature-level holdout validation, all perturbed transcriptomic signatures in the test set were unseen. Each signature represents drug-induced gene expression change for all 19216 genes under a certain combination of drug treatment, time duration, dosage, and cellular context. For drug-level holdout validation, all drugs in the test set were unseen. F1 score and balanced accuracy between prediction and ground truth are used as evaluation metrics. All the box-plots show the study-wise performance. n represents the number of signatures in the test set. **B,** Scatter plots showing correlation between signature-wise prediction performance and estimated perturbation magnitude score (Methods). n represents the number of perturbed transcriptomic signatures in the test set. **C,** Boxplots showing performance between all signatures and signatures with high perturbation magnitude scores of >=0.3 in the test set. n represents the number of signatures in the test set. **D,** Boxplots showing performance on the test sets between using all signatures and using strong-signal signatures for fine-tuning / testing. Strong-signal signatures are whose label variance of predicted expression across the transcriptome >0.1 or >0.35 (Methods). n represents the number of signatures in the test sets. Asterisks mean statistically significant difference via t-test (*: < 0.05, ** < 0.01, *** < 0.001).

**Figure S7.**
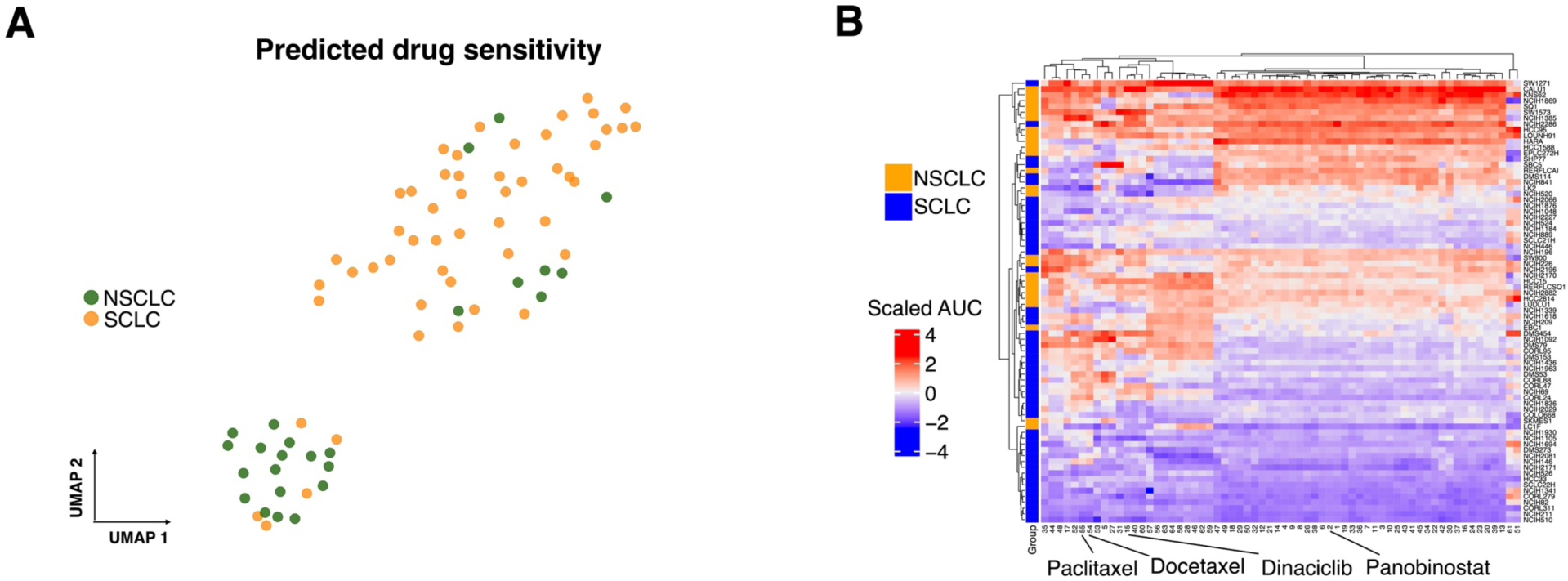
Additional results on using InsilicoCell for phenotype-based drug screening in lung cancer, related to Figure 3. **A,** Visualization of the drug sensitivity predicted by InsilicoCell for all 816,230 screened drugs in NSCLC and SCLC cell lines. Compounds and cell lines were all collected from combined LINCS, BindingDB and CTRP databases. Two distinct clusters revealed that InsilicoCell captured disease subtype-specific differences in drug sensitivity. **B**, Heatmap showing the drug sensitivity predicted by InsilicoCell for the top drug candidates in NSCLC and SCLC cell lines. Several drugs with previously reported efficacy in lung cancer treatment are labeled. InsilicoCell identified FDA-approved treatments such as docetaxel and paclitaxel, to be among the top 50 potent candidates for NSCLC from the initial large pool of 816,230 compounds. It also showed some other candidates to be slightly more specific for SCLC compared to NSCLC, such as panobinostat.

**Figure S8.**
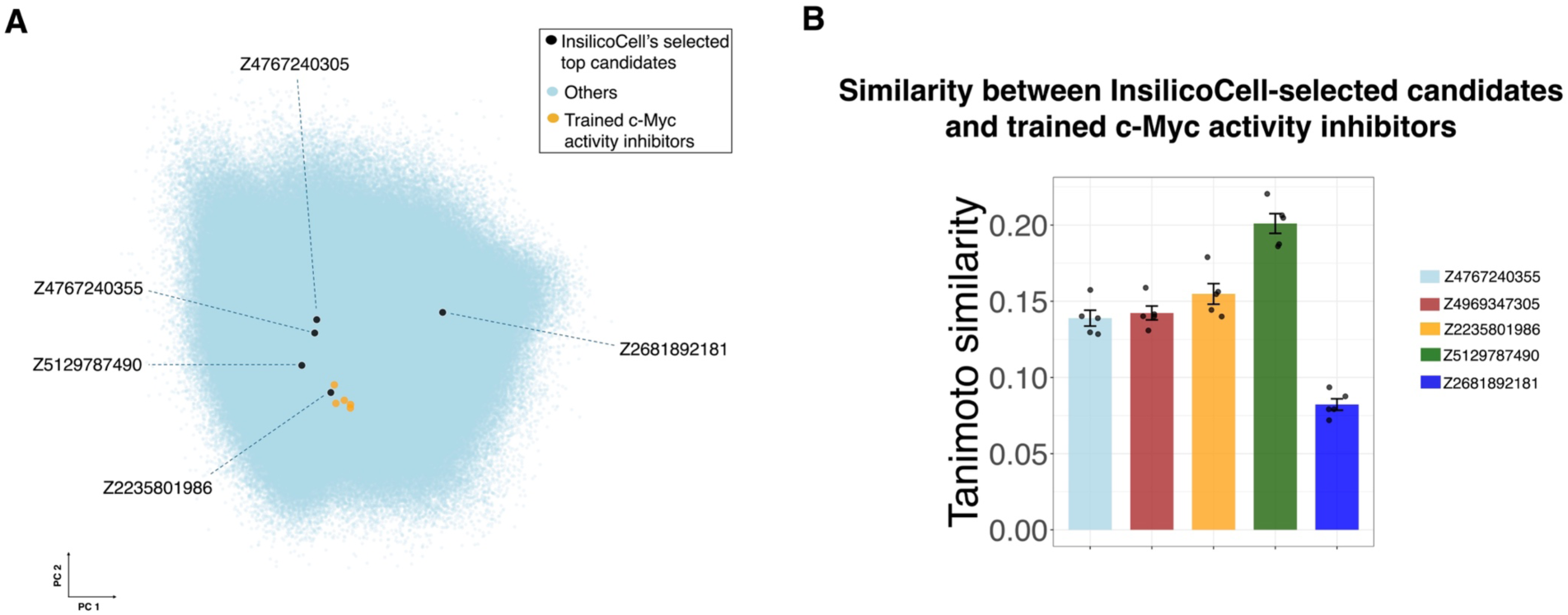
Novelty analysis on the c-Myc activity inhibitors predicted by InsilicoCell in HCC, related to Figure 4. **A,** Chemical structure space of library compounds, highlighting the final selected candidates: Z5129787490, Z4767240355, Z4767240305, Z2235801986, Z2681892181 using black color, as well as the c-Myc activity inhibitors from pretrained data using orange color. **B,** Tanimoto similarity between each InsilicoCell-predicted candidate and each pretrained c-Myc activity inhibitor, based on ECFP representations of the compounds.

**Figure S9.**
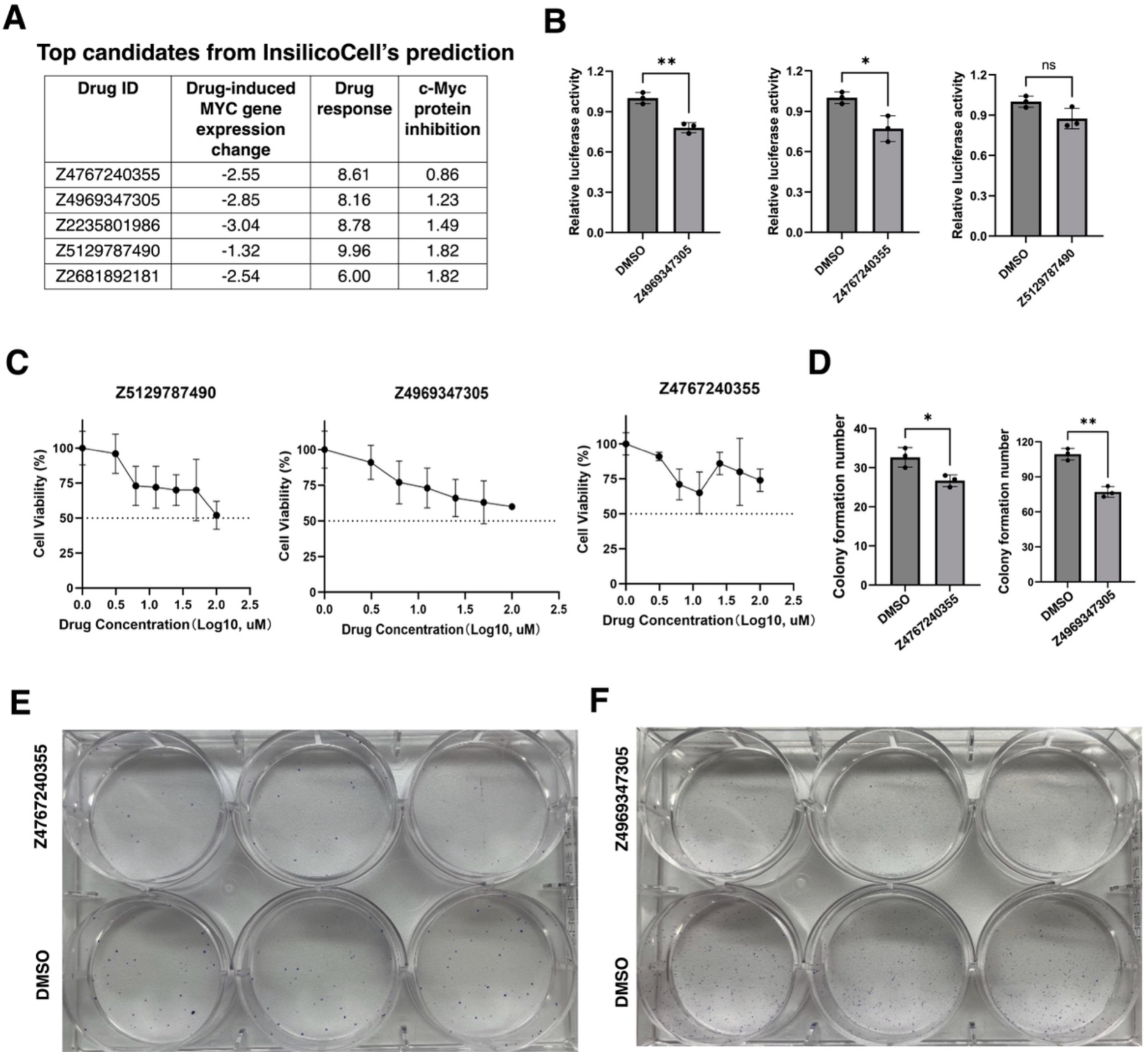
Additional results on using InsilicoCell for multi-objective screening for novel cMyc activity inhibitors in HCC, related to Figure 4. **A,** Drug candidates selected by InsilicoCell and their prediction values for the three objectives. **B,** Dual-luciferase reporter assay showing cMyc protein activity inhibition with Z4767240305, Z4767240355, and Z5129787490. **C**, Dose-response curves of Z4767240305, Z4767240355, and Z5129787490 in HepG2 cells. **D-F**, Colony formation results showing inhibition of HepG2 cell growth under treatment of Z4767240355 and Z4767240305 for 14 days. Asterisks mean statistically significant difference via t-test (*: < 0.05, ** < 0.01, *** < 0.001).

**Figure S10.**
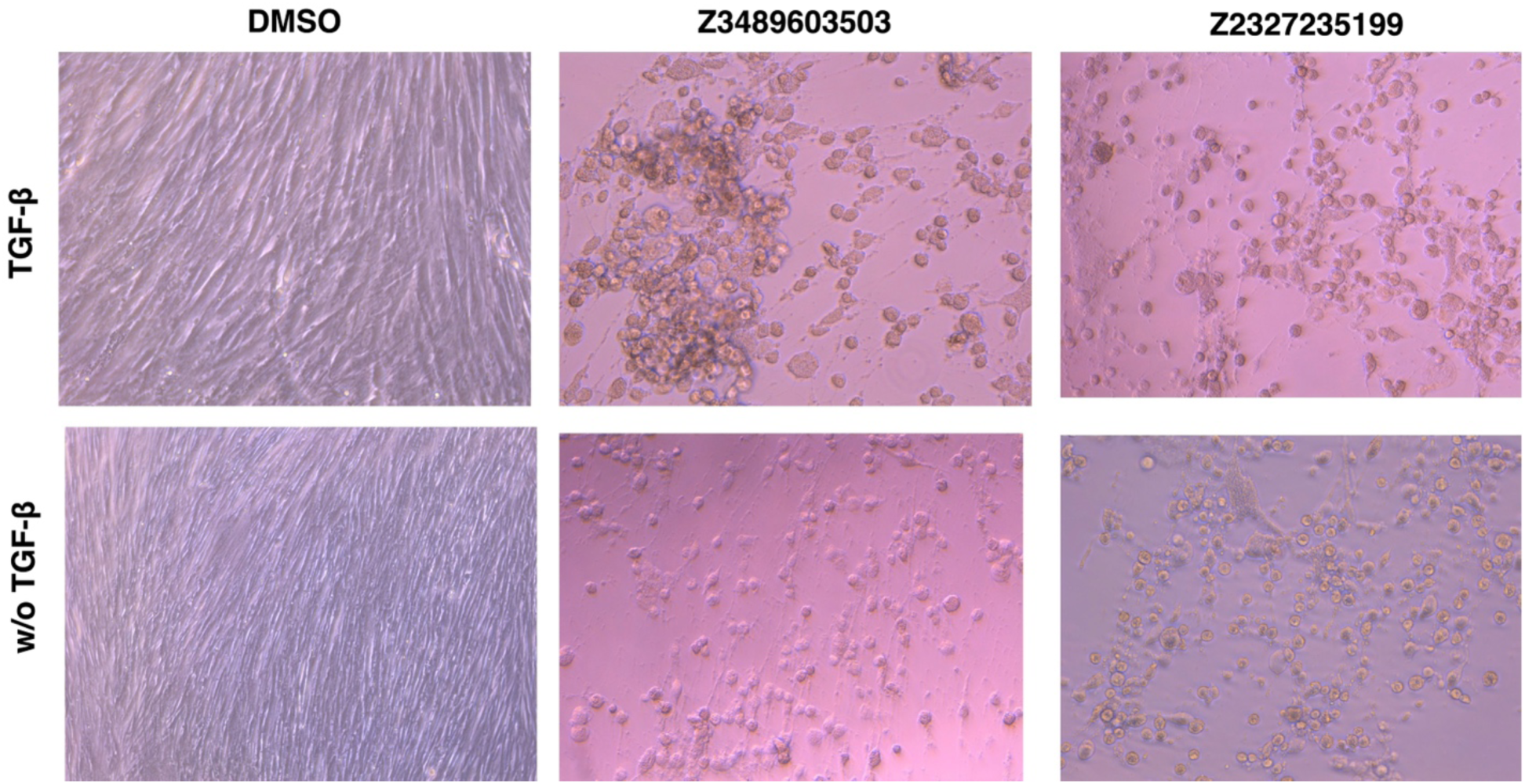
Additional results on using InsilicoCell for multi-objective screening for drugs inhibiting fibrosis markers in myofibroblasts, related to Figure 5. Microscopy showing cell death of myofibroblasts (TGF-β) and fibroblasts (w/o TGF-β) treated with 10μM Z3489603503 or Z2327235199 for 3 days.

**Figure S11.**
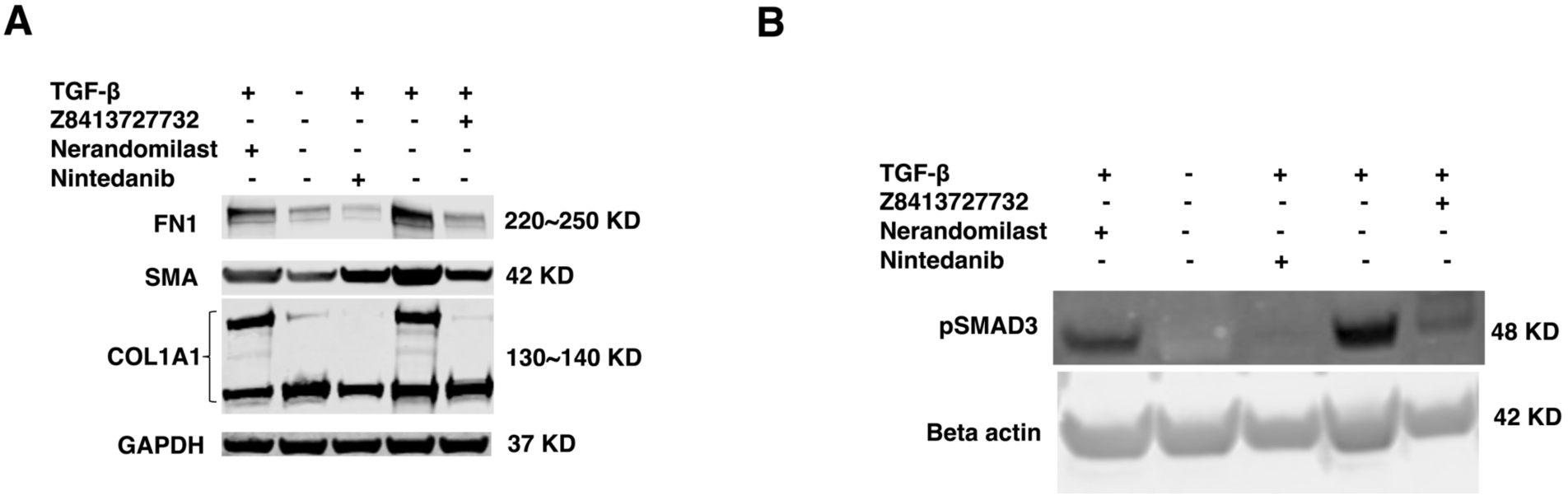
Additional western-blot results on using InsilicoCell for multi-objective screening for drugs inhibiting fibrosis markers in IPF, related to Figure 5. **A**, Western blot showing reduced expression of fibrosis markers COL1A1, SMA, and FN1 in TGF-*β* induced myofibroblasts under treatment of 10μM Z8413727732 compared to two FDA-approved IPF drugs, nerandomilast and nintedanib. **B**, Western-blot results showing inhibition of pSMAD3 in TGF-*β* induced myofibroblasts under treatment of 10μM Z8413727732, compared to two FDA-approved IPF drugs, nerandomilast and nintedanib.

**Figure S12.**
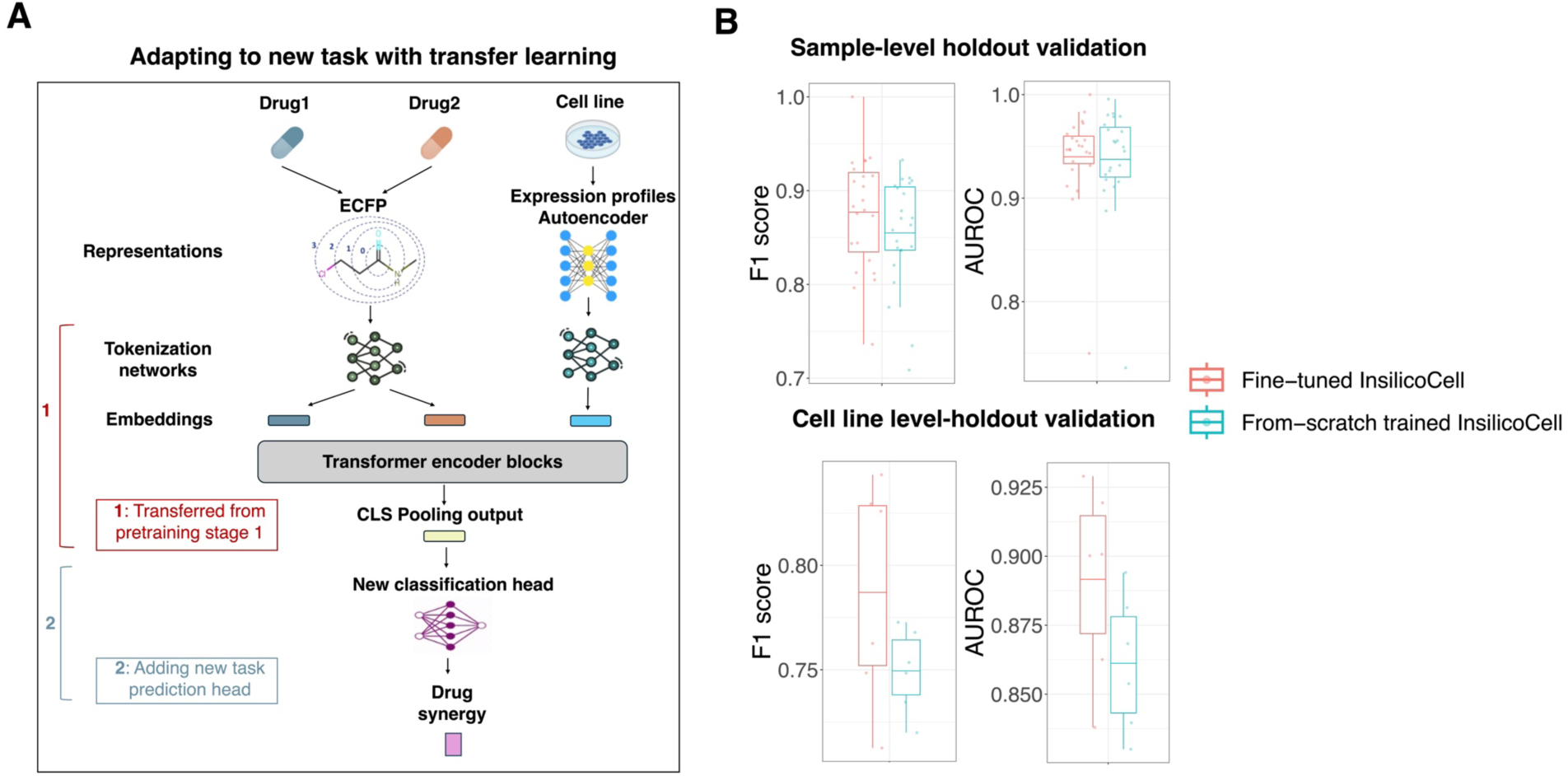
Applying InsilicoCell to drug synergy prediction via transfer learning, related to Figure 7A, Diagram showing the transfer learning approach for adapting the pretrained InsilicoCell to the new binary classification task of drug combination synergy prediction. **B**, Comparison of the prediction performance between fine-tuned InsilicoCell, which was pretrained on the seven types of cellular functional profiles shown in Figure 1A, and the from-scratch trained InsilicoCell, which was not pretrained on the seven types of cellular functional profiles. In the boxplots, one dot represents one cell line.

**Table S1.** Summary of data sets used for pretraining and testing*.

| Name | Data type | # of contexts (cell lines / patients / single cells / spatial spots) | # of drugs (compounds) | # of proteins / TFs | # of genes | # of time points | # of dosages | # of labeled samples | Usage |
| --- | --- | --- | --- | --- | --- | --- | --- | --- | --- |
| LINCS (L5) | Drug-induced gene expression change | 82 | 5768 |  | 964 | 5 | 6 | 29459840 | pretrain & test splitting |
| Binding DB | Drug-protein binding affinity* |  | 810607 | 6416 |  |  |  | 1549276 | pretrain & test splitting |
| CTRP | Drug sensitivity | 826 | 531 |  |  |  |  | 339120 | pretrain & test splitting |
| DepMap | Gene effect score | 621 |  |  | 17479 |  |  | 10824961 | pretrain & test splitting |
| ChEA | Transcription factor (TF)-target gene association |  |  | 199 | 17344 |  |  | 3451456 | pretrain & test splitting |
| DepMap | Gene mutation | 1203 |  |  | 17137 |  |  | 20615811 | pretrain & test splitting |
| DepMap | Copy number variation (CNV) | 1196 |  |  | 18333 |  |  | 21926268 | pretrain & test splitting |
| Bottomly D et al., 2022 | Drug sensitivity | 394 | 141 |  |  |  |  | 42413 | patient-level test & fine-tuning splitting |
| Wu SZ et al., 2021 | Spatial transcriptome | 2426 (ER+) / 4895 (TNBC) |  |  |  |  |  |  | spatial-level test |
| Habermann AC et al., 2020 | Single-cell transcriptome | 114396 |  |  |  |  |  |  | IPF; single cell-level test |
| GDSC | Drug sensitivity | 281 | 29 |  |  |  |  | 4727 | External validation |
| ChEMBL | Drug-protein binding affinity* |  | 20715 | 232 |  |  |  | 22507 | External validation |
| GEOMeta | Drug-induced gene expression change | 953 | 813 |  | 19216 | 11 | 14 | 50422784 | External validation & fine-tuning splitting |
| GDSC<sup>2</sup> | Drug combination synergy | 31 | 38 |  |  |  |  | 13243 | New task |

\* All data sets listed were after preprocessing, where the numbers of contexts, drugs, proteins, genes, time points and dosages represent the actual numbers of these entities for model training and testing. They may differ from the original numbers of entities in the raw data.
\* Most drug-protein binding affinities represent direct physical ligand-protein interactions. However, measurements from functional or inhibition assays may not necessarily indicate direct binding, as exemplified by the c-Myc activity data in this case study.

**Table S2.** Top 50 compound candidates selected by InsilicoCell in lung cancer cell lines. 36 candidates were shared between NSCLC and SCLC.

| Drug index | Rank in NSCLC | Rank in SCLC | Predicted AUC in NSCLC | Predicted AUC in SCLC | Drug name / PubChem CID | MOA |
| --- | --- | --- | --- | --- | --- | --- |
| 1 | 4 | 1 | 5.82 | 3.56 | 139558701 | HDAC inhibitor |
| 2 | 3 | 2 | 5.81 | 3.56 | Panobinostat | HDAC inhibitor |
| 3 | 6 | 3 | 6.16 | 3.88 | 155517175 | HDAC inhibitor |
| 4 | 10 | 4 | 6.44 | 3.93 | 53363889 | HDAC inhibitor |
| 5 | 1 | 5 | 4.31 | 3.94 | Leptomycin B | CRM1 inhibitor |
| 6 | 9 | 6 | 6.35 | 3.99 | 137647813 | HDAC inhibitor |
| 7 | 7 | 7 | 6.2 | 3.99 | 155526247 | HDAC inhibitor |
| 8 | 11 | 8 | 6.44 | 4.11 | 20579392 | HDAC inhibitor |
| 9 | 15 | 9 | 6.69 | 4.14 | 20579393 | HDAC inhibitor |
| 10 | 14 | 10 | 6.62 | 4.16 | 155525662 | HDAC inhibitor |
| 11 | 12 | 11 | 6.44 | 4.2 | 155537455 | HDAC inhibitor |
| 12 | 17 | 12 | 6.79 | 4.37 | 20579374 | HDAC inhibitor |
| 13 | 26 | 13 | 7.03 | 4.42 | 53363970 | HDAC inhibitor |
| 14 | 19 | 14 | 6.86 | 4.42 | 20579375 | HDAC inhibitor |
| 15 | 5 | 15 | 6.13 | 4.54 | Dinaciclib | CDK inhibitor |
| 16 | 32 | 16 | 7.21 | 4.54 | 155552802 | HDAC inhibitor |
| 17 | 8 | 17 | 6.3 | 4.72 | SB-743921 | KSP inhibitor |
| 18 | 30 | 18 | 7.16 | 4.73 | 44205087 | HDAC inhibitor |
| 19 | 45 | 19 | 7.42 | 4.76 | 139377827 | HDAC inhibitor |
| 20 | 47 | 20 | 7.47 | 4.86 | 118712705 | HDAC inhibitor |
| 21 | 31 | 21 | 7.2 | 4.87 | 11024529 | HDAC inhibitor |
| 22 | 24 | 22 | 6.99 | 4.9 | 53363971 | HDAC inhibitor |
| 23 | 49 | 23 | 7.59 | 4.94 | 155539714 | HDAC inhibitor |
| 24 | 43 | 24 | 7.37 | 4.94 | 155537405 | HDAC inhibitor |
| 25 | 25 | 25 | 7.01 | 4.98 | 53363805 | HDAC inhibitor |
| 26 | 52 | 26 | 7.62 | 5 | 9858986 | HDAC inhibitor |
| 27 | 2 | 27 | 5.69 | 5.03 | 45258492 | CRM1 inhibitor |
| 28 | 18 | 28 | 6.85 | 5.12 | ouabain | Na<sup>+</sup>/K<sup>+</sup>-ATPase inhibitor |
| 29 | 50 | 29 | 7.6 | 5.12 | 53364355 | HDAC inhibitor |
| 30 | 27 | 30 | 7.09 | 5.17 | BJB-432 | HDAC inhibitor |
| 31 | 16 | 31 | 6.71 | 5.17 | CR-1-31-B | eIF4A inhibitor |
| 32 | 28 | 32 | 7.13 | 5.2 | N/A | Unknown |
| 33 | 40 | 33 | 7.37 | 5.22 | N/A | Unknown |
| 34 | 60 | 34 | 7.78 | 5.23 | 9880847 | HDAC inhibitor |
| 35 | 54 | 35 | 7.66 | 5.24 | GSK461364 | PLK1 inhibitor |
| 36 | 77 | 36 | 7.94 | 5.27 | 139368356 | HDAC inhibitor |
| 37 | 56 | 37 | 7.68 | 5.27 | 155522601 | HDAC inhibitor |
| 38 | 55 | 38 | 7.67 | 5.32 | 10894419 | HDAC inhibitor; in lung cancer |
| 39 | 59 | 39 | 7.75 | 5.34 | 24959169 | HDAC inhibitor |
| 40 | 23 | 40 | 6.99 | 5.35 | 145989210 | CDK inhibitor |
| 41 | 44 | 41 | 7.39 | 5.38 | 53363887 | HDAC inhibitor |
| 42 | 57 | 42 | 7.69 | 5.38 | 118712713 | Unknown |
| 43 | 36 | 43 | 7.31 | 5.4 | 53363806 | HDAC inhibitor |
| 44 | 116 | 44 | 8.31 | 5.44 | LSM-1797 | Unknown |
| 45 | 114 | 45 | 8.3 | 5.47 | 118712701 | HDAC inhibitor |
| 46 | 34 | 46 | 7.27 | 5.47 | Dihydroouabain | Na<sup>+</sup>/K<sup>+</sup>-ATPase inhibitor |
| 47 | 73 | 47 | 7.91 | 5.47 | 44233027 | HDAC inhibitor |
| 48 | 123 | 48 | 8.34 | 5.5 | 6324671 | Unknown |
| 49 | 94 | 49 | 8.12 | 5.51 | 155561265 | HDAC inhibitor |
| 50 | 100 | 50 | 8.21 | 5.52 | PD083355 | Unknown |
| 51 | 13 | 134 | 6.46 | 6.49 | 72708130 | Opioid receptor |
| 52 | 20 | 54 | 6.9 | 5.61 | N/A | Unknown |
| 53 | 21 | 78 | 6.93 | 5.97 | 137650531 | CRM1 inhibitor |
| 54 | 22 | 96 | 6.96 | 6.1 | docetaxel | FDA-approved lung cancer drug; NSCLC; tubulin modulator |
| 55 | 29 | 62 | 7.15 | 5.78 | paclitaxel | FDA-approved lung cancer drug; NSCLC; tubulin modulator |
| 56 | 33 | 61 | 7.22 | 5.75 | 137641025 | Na<sup>+</sup>/K<sup>+</sup>-ATPase inhibitor |
| 57 | 35 | 149 | 7.29 | 6.56 | Omacetaxine<br>Mepesuccinate | Unknown |
| 58 | 37 | 83 | 7.31 | 6.01 | 9828429 | Unknown |
| 59 | 38 | 51 | 7.33 | 5.57 | digoxin | Na<sup>+</sup>/K<sup>+</sup>-ATPase inhibitor |
| 60 | 39 | 53 | 7.35 | 5.6 | 168269482 | CDK inhibitor |
| 61 | 41 | 489 | 7.37 | 7.96 | 72708197 | Opioid receptor |
| 62 | 42 | 55 | 7.37 | 5.61 | digitoxin | Na<sup>+</sup>/K<sup>+</sup>-ATPase inhibitor |
| 63 | 46 | 72 | 7.46 | 5.89 | 137644838 | Na<sup>+</sup>/K<sup>+</sup>-ATPase inhibitor |
| 64 | 48 | 102 | 7.52 | 6.14 | 101398153 | Na<sup>+</sup>/K<sup>+</sup>-ATPase inhibitor |

**Table S3.** *MYC* gene effect scores predicted by InsilicoCell in HCC cell lines. A negative score below −1 indicates the corresponding cell line is highly dependent on MYC for survival.

| Cell line | Predicted *MYC* gene effect score |
| --- | --- |
| SNU878 | -2.41 |
| JHH6 | -2.07 |
| JHH5 | -2.05 |
| SNU398 | -2.03 |
| SNU387 | -2.01 |
| JHH1 | -1.93 |
| SNU475 | -1.89 |
| JHH4 | -1.88 |
| PLCPRF5 | -1.86 |
| HEPG2 | -1.85 |
| HUH7 | -1.85 |
| SNU449 | -1.77 |
| JHH7 | -1.75 |
| HEP3B217 | -1.7 |
| SNU182 | -1.69 |
| SNU423 | -1.57 |
| HLF | -1.53 |

## Notes

### Competing Interest Statement

The authors have declared no competing interest.

